# Dopamine modulation of prefrontal cortex activity is manifold and operates at multiple temporal and spatial scales

**DOI:** 10.1101/452862

**Authors:** Sweyta Lohani, Adria K. Martig, Karl Deisseroth, Ilana B. Witten, Bita Moghaddam

**Author notes:** Address correspondence to: Bita Moghaddam, Department of Behavioral Neuroscience, Oregon Health & Sciences University, Mail code L-470 3181 S.W. Sam Jackson Park Rd, Portland, OR 97239-3098.

## Abstract

While the function of dopamine in subcortical structures is largely limited to reward and movement, dopamine neurotransmission in the prefrontal cortex (PFC) is critical to a multitude of temporally and functionally diverse processes such as attention, working memory, behavioral flexibility, action selection, and stress adaptation. How does dopamine influence PFC computation of multiple temporally diverse functions? Here we find causation between sustained and burst patterns of phasic dopamine neuron activation and contemporaneous modulation of PFC neuronal activity at multiple spatio-temporal scales. These include a multidirectional and weak impact on individual PFC neuron rate activity and a robust influence on coordinated ensemble activity, gamma oscillations, and gamma-theta coupling that persisted for minutes. In addition, PFC network responses to burst pattern of dopamine firing were selectively strengthened in behaviorally active states. Thus, dopamine modulation of PFC is spatiotemporally diverse and is dictated by the pattern of dopamine neuron activation and behavioral state. These findings provide insight on the multiplex pattern of modulation by dopamine that may influence PFC computation of temporally diverse functions.

## Introduction

The pioneering work of Patricia Goldman-Rakic ^1^ and Anne-Marie Thierry ^2^ described two novel roles for dopamine in the prefrontal cortex (PFC) that were dynamically and functionally distinct from its previously described subcortical roles in mediating reward ^3^ and movement ^4^. Specifically, dopamine neurons localized in the ventral tegmental area (VTA) that projected to the PFC were uniquely sensitive to acute stress ^2^, and PFC dopamine was shown to be necessary for working memory function ^1^. A large body of basic and clinical research has since established that in addition to influencing stress-related function and working memory, dopamine neurotransmission in PFC plays a role in nearly all aspects of high order cognition, including attention and behavioral flexibility ^5, 6^; PFC dopamine is also implicated in cognitive deficits of brain illnesses such as schizophrenia, addiction, and ADHD ^7, 8^. An equally large literature has expanded on PFC dopamine’s role in stress response and adaptation as well as stress or anxiety related disruption of goal-directed behavior, including working memory ^9^.

Whereas much advance has been made in understanding how dopamine neurotransmission computes reward ^10, 11^ and action ^12-14^, our understanding of how dopamine broadcasts its multifaceted effect on PFC neuronal activity remains poorly understood. A challenge has been that the functions attributed to PFC dopamine are temporally variable (e.g. ms-sec for attentional tuning, 1-40 sec for working memory, and sec-min for stress or anxiety modulation of behavior) and thus likely involve modulation of both persistent and dynamic activity of PFC neurons. Another challenge is that dopamine transporter density is considerably lower in PFC compared to striatal regions ^15^, and dopamine receptors in PFC are mostly distant from release sites ^16^. Thus, dopamine can travel farther in the PFC compared to subcortical structures, making the post-release impact of dopamine spatially and temporally more complex than that in striatal regions. In fact, previous reports which have studied dopamine’s action on PFC activity as direct input-output linear relationships—that is, assuming that an increase in dopamine input causes an increase or decrease on the firing rate of individual PFC neurons—have produced mixed and altogether inconsistent results (e.g.^17-20^).

We wanted to better understand how changes in the activity of dopamine neurons affect the neurodynamic properties of PFC. Based on previous findings, we postulated that the input-output impact of dopamine on PFC activity is not a linear rate (increase or decrease) effect. Instead, information propagation by dopamine in the PFC may involve multiple spatiotemporal output responses. To investigate this, we made concomitant assessment of individual neuronal spiking, population ensemble activity, and local field potential oscillations as we stimulated dopamine neurons in the VTA using optogenetics in *Th::Cre* rats ^21^. Experiments were performed during wake cycle after animals were habituated to the recording chamber to, as much as possible, reduce the impact of stress and cognitive load on spontaneous dopamine or PFC neuron activity. Dopamine neurons were stimulated using protocols mimicking three different activity patterns: slow and fast phasic burst activity that are generally associated with reward-prediction or instructive signaling ^22^, and sustained phasic activity that has been observed during tasks involving sustained attention and distant-goals ^23, 24^. We examined both transient and prolonged effects of these different patterns of phasic activation of dopamine neurons on PFC neural activity in spontaneously behaving animals.

## Results

### Optogenetic stimulation entrained VTA dopamine neurons

We injected AAV5-DIO-ChR2-eYFP into the VTA to target dopamine neurons in *Th::Cre* rats (Figure 1A) and in wild-type littermates to control for non-specific effects of stimulation. All studies were performed at least four weeks after virus injection to allow for sufficient integration. We have previously verified highly specific and sensitive expression of ChR2 in VTA dopamine neurons of these *Th::Cre* rats ^25^. Here, we established that dopamine neurons can entrain to different patterns of optogenetic stimulation. VTA dopamine neurons were stimulated and simultaneously recoded from via an optrode (combined electrode and optical fiber) during the wake cycle of *Th::Cre* rats in a recording chamber (Figure 1B). We used different stimulation parameters that were similar to patterns of dopamine cell firing during active behavior (Figure 1C). Specifically, dopamine neurons can fire in a sustained phasic pattern and, more classically, in bursts with intra-burst frequencies between 15 – 100 Hz. The sustained phasic pattern of firing has been reported when an upcoming cue needs to be detected during tasks that involve distant goals and sustained attention ^23^. An example of a raw trace of this response pattern is shown on the top panel of Figure 1D. To model the pattern of “sustained phasic activation”, we delivered a train of blue light pulses (20 Hz, train width = 5 s) every 10 s. The second pattern involving burst firing is widely reported in response to various salient events such as cues, novel/unexpected rewards, and absence of expected rewards ^22, 26, 27^. An example of VTA dopamine neuron burst firing in response to reward is shown on the bottom panel of Figure 1D. We used two protocols to model burst activation: 100 Hz “fast burst” and 20 Hz “slow burst”. Bursts were repeated every 500 ms (Figure 1C). All stimulation parameters were applied for 10-20 min because maze or operant asks typically last for this duration during which multiple trials (approximately 100 for fixed ratio operant tasks) are repeated ^23, 28, 29^.

**Figure 1.**
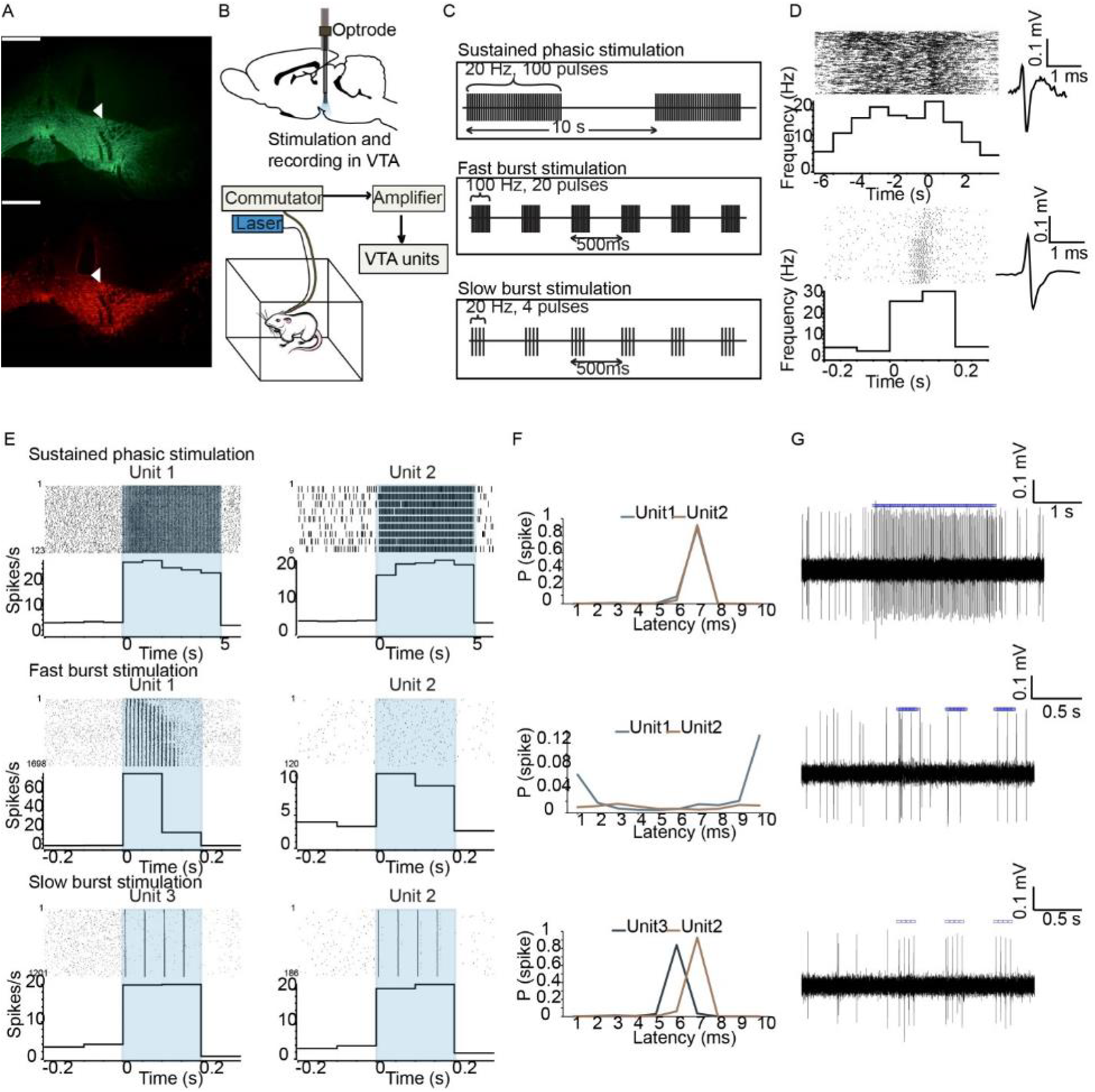
**A.** Expression of ChR2-eYFP (top) and TH (bottom) in the VTA of a representative *Th::Cre* rat. Scale bar = 550 µm. White triangles point to the termination of an optical fiber implant just dorsal to VTA. **B.** (Top) Schematic of implantation of an optrode (combined optical fiber and electrode) in the VTA. (Bottom) Simultaneous optogenetic stimulation and recording in VTA in freely moving *Th::Cre* rats. **C.** Illustrations of stimulation protocols. **D.** Example phasic responses of VTA dopamine neurons to behavioral events (left) and the corresponding neuron’s waveform (right). (Top) VTA dopamine neuron showed a sustained phasic response prior to cue onset (t = 0 s, bin size = 1 s) during an attention task, adapted from ^23^. (Bottom) VTA dopamine neuron firing exhibited a transient burst during reward delivery (t = 0 s, bin size = 0.1 s) in a Pavlovian conditioning task, adapted from ^74^. **E.** Three well-isolated VTA units (from n = 2 *Th::Cre* rats) responded to VTA optogenetic stimulation. Responses of units to different stimulation protocols are presented in separate rows as rasters (top) and peri-event histograms (bottom; bin size = 1 s (sustained phasic), bin size = 0.1 s (fast and slow burst)). On rasters, trial numbers are shown on the left (note: total trial numbers are different for each raster). T = 0 indicates start of a burst or phasic train and blue rectangles mark the period of stimulation. **F.** Probability of unit spiking at latencies 0-10 ms from the delivery of single blue light pulses. Each panel corresponds to the rasters/histograms presented in the same row in 1E (See Supplementary Figure 1 for comparison of spike probabilities in early versus late trials). **G.** Raw voltage traces for unit 2 showing responses to sustained phasic (top), fast burst (middle), and slow burst (bottom) stimulation. Blue squares indicate when each blue light pulse was delivered.

Responses of three well-isolated example VTA units to different stimulation protocols are shown in Figure 1E. These units responded reliably to light pulses in the sustained phasic stimulation and slow burst stimulation protocols (probability of spiking with a latency of < 10 ms was > 90%) (Figure 1F), while units responded with lower fidelity to the fast burst stimulation pulses (probability of spiking for unit 1 at a latency of < 10 ms was 28% and for unit 2 was 10 %) (Figure 1F). Despite the overall low probability of spiking during fast burst stimulation, Unit 1 showed more reliable responses to pulses presented earlier in the burst compared to later in the burst (Figure 1E). Furthermore, from observing the rasters and histograms (Figures 1E, 1F) as well as raw voltage traces (Figure 1G) for these units, it is apparent that fast burst stimulation elicited transient bursts with a few spikes emitted at a frequency of 10 – 100 Hz. Thus, units respond to fast burst stimulation phasically but with large variability as indicated by variable inter-spike-intervals (ISIs) and number of spikes per burst. (Based on our experience of recording from VTA neurons in behaving rodents, these response patterns mimic naturalistic firing patterns in dopamine neurons as a given task-responsive neuron does not typically fire with fixed ISIs or with fixed burst spike number). For all stimulation protocols, there was no apparent change in light responsivity of dopamine neurons over time due to factors such as ChR2 desensitization or depolarization block, as their probabilities of spiking in response to light pulses were similar in early and late trials (Supplementary Figure 1).

### Optogenetic stimulation of VTA dopamine neurons increased mPFC extracellular dopamine levels

To ensure that both sustained phasic and burst optogenetic activation of VTA dopamine cells release dopamine in mPFC, PFC microdialysis was conducted. A separate cohort of *Th::Cre* and wild-type rats that were infused with the AAV viral construct in VTA was implanted with microdialysis probes in mPFC and optical fibers in VTA (Figure 2A and Supplementary Figure 2). We found that VTA optogenetic stimulation with the sustained phasic and fast burst protocols increased mPFC extracellular dopamine levels ([DA]_o_) on average by >200% of baseline selectively in *Th::Cre* but not wild-type rats. Sustained phasic and fast burst protocols similarly increased mPFC [DA]_o_ (two way repeated measures ANOVA, stim: F(1,5) = 0.174, p = 0.694; sample X stim type, F(2,4) = 7.983, p = 0.492). The increase in [DA]_o_ was sustained during post-stimulation samples (Figure 2B and 2C). The magnitude and pattern of dopamine increases were similar to previously reported increases in [DA]_o_ when animals are performing a reward-driven behavioral flexibility task ^28^, suggesting that our stimulation protocols elicited physiologically relevant sustained dopamine release in mPFC.

**Figure 2.**
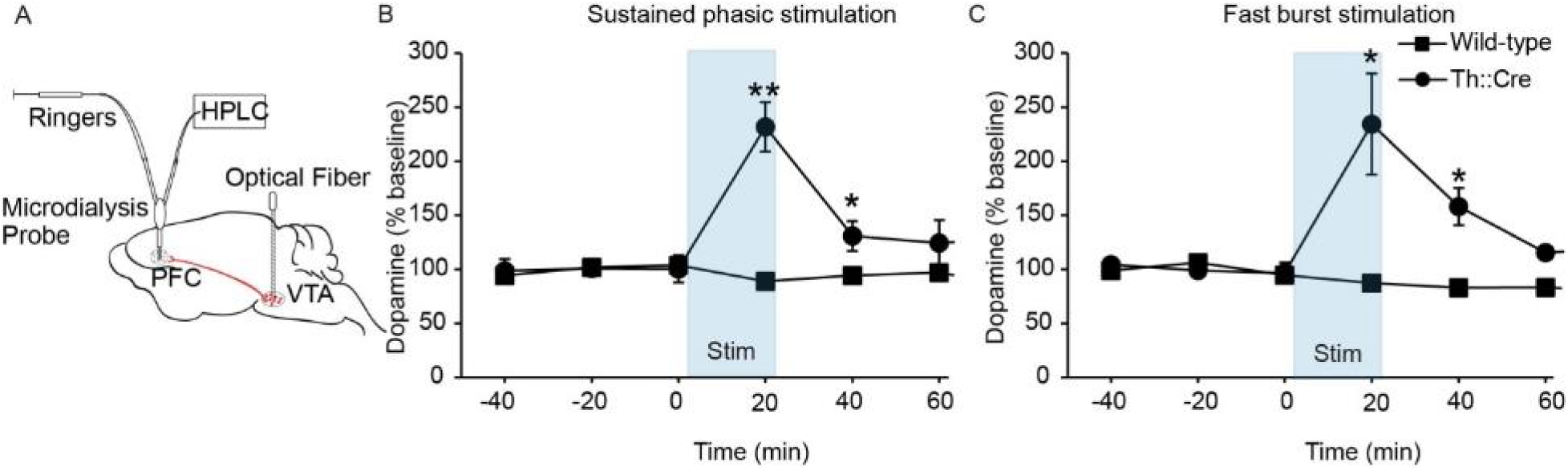
Sustained and burst phasic activation of VTA dopamine neurons increased [DA]_o_ in mPFC. **A**. Schematic of microdialysis setup. **B.** Sustained phasic stimulation of VTA dopamine neurons for 20 min significantly increased mPFC [DA]_o_ (F(8,24) = 6.536, p = 0.001, repeated measures ANOVA) in stimulation sample 4 and post-stimulation sample 5 compared to baseline (sample 4: p = 0.002, sample 5: p = 0.043; t-tests) in *Th::Cre* rats (n = 4 rats, mean baseline [DA]_o_ = 0.114 +/− 0.088 fmol). Sustained phasic stimulation in wild-type rats (n = 3 rats, mean baseline [DA]_o_ = 0.136 +/− 0.026 fmol) did not increase [DA]_o_ (F(8,16) = 1.016, p = 0.462). **C.** Fast burst stimulation of VTA dopamine neurons for 20 min significantly increased [DA]_o_ (F (8,16) = 6.107, p = 0.005) in stimulation sample 4 and post-stimulation sample 5 (sample 4: p = 0.026, sample 5: p = 0.026; t-tests) in *Th::Cre* rats (n = 3 rats, mean baseline [DA]_o_ = 0.109 +/− 0.052 fmol). Wild-type rats (n = 3 rats, mean baseline [DA]_o_ = 0.220 +/− 0.108 fmol) did not show an increase in [DA]_o_ after fast burst VTA stimulation (F(8,16) = 2.231, p = 0.228). Furthermore, two way repeated measures ANOVAs indicated that the pattern of [DA]_o_ change over time was significantly different between *Th::Cre* and wild-type rats after stimulation (sustained phasic stimulation: F(8,40) = 4.362, p = 0.001, fast burst stimulation: F(8,32) = 6.075, p = 0.000; interaction effects). Error bars indicate mean +/− SEM. ** p < 0.05 and * p < 0.01. (See Supplementary Figure 2 for histological confirmation of placements).

### Activation of dopamine neurons elicited a heterogeneous response on multiple timescales in a minority of PFC units

Next, we examined the impact of sustained phasic and burst activation of dopamine neurons on PFC neural activity (Figure 3A,B & Supplementary Figure 3). During a recording session, only one of three stimulation protocols (illustrated in Figure 1C) was utilized, and the order of stimulation protocols was counterbalanced across sessions that were separated by at least 24 hours. Fast and slow burst stimulation lasted for 10 min, during which the total time of illumination (laser on) was identical for the two protocols (= 24 s). Fast and slow burst protocols resulted in ~1200 bursts or transient trials during the 10 min of stimulation. Sustained phasic stimulation, on the other hand, resulted in only 60 phasic trains/transient trials during 10 min of stimulation even though the total amount of light exposure was comparable (total time of illumination in 10 min = 30 s). To increase the number of transient trials, sustained phasic stimulation was continued for 20 min.

**Figure 3.**
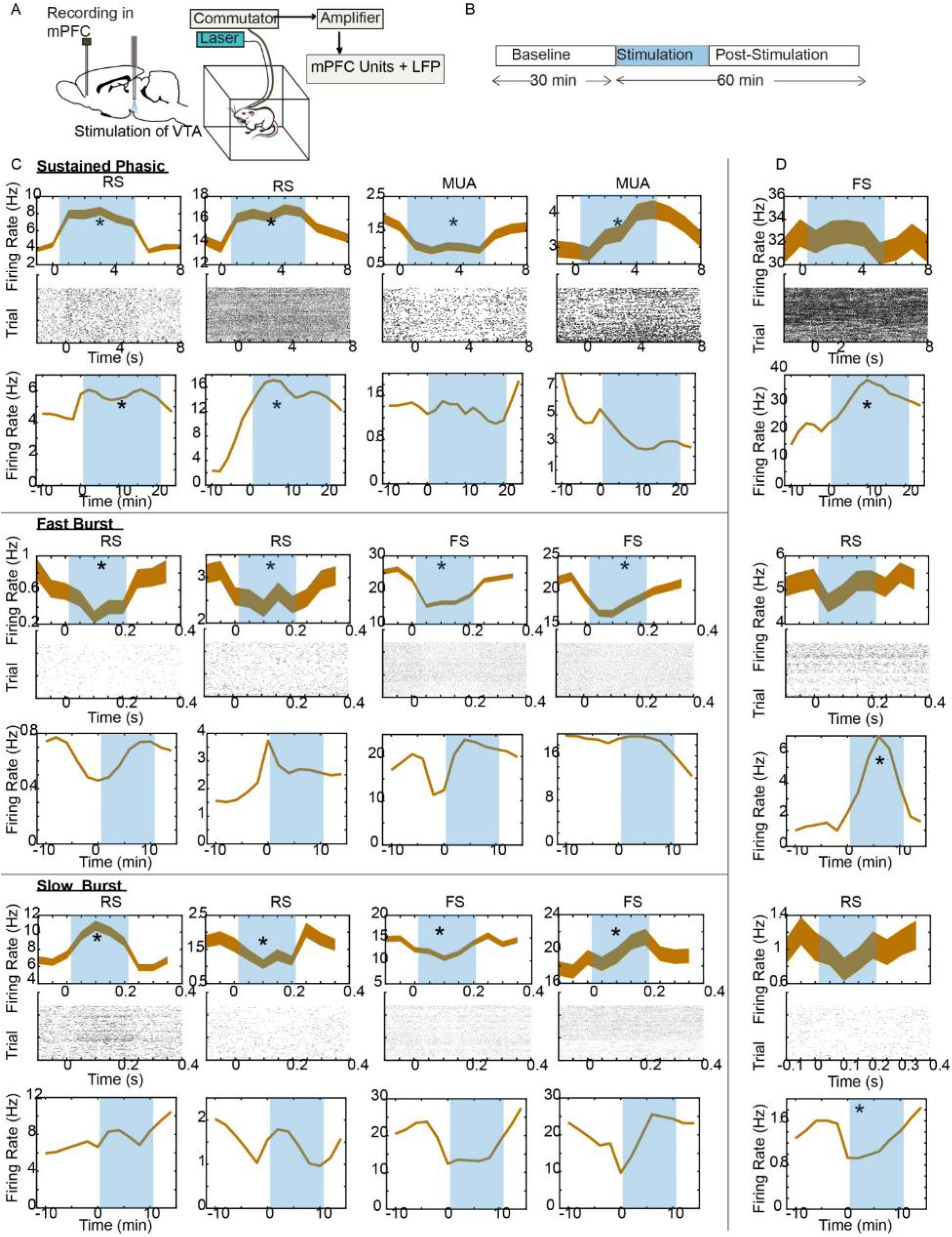
Recording schematic and example unit responses. **A.** Schematic of simultaneous optogenetic stimulation of VTA and recording of LFPs and unit activity in mPFC of freely moving rats. **B.** Recording timeline in a session. **C.** Example units in mPFC that are significantly modulated on a transient timescale but not necessarily on a prolonged timescale. Responses of example regular-spiking (RS) single units, fast-spiking (FS) single units or multi-units (MUA) in mPFC to optogenetic stimulation of VTA dopamine neurons with sustained phasic, fast burst, and slow burst protocols are illustrated. For each stimulation protocol, the top two rows show transient responses (on the order of seconds) of units as peri-event firing rate histograms (mean +/− SEM, bin size = 1s for sustained phasic protocol and bin size = 0.05 s for fast and slow burst protocols) and rasters. The bottom row for each protocol shows prolonged responses (on the order of minutes) as peri-event firing rate histograms (bin size = 120 s) to repeated phasic stimulation of VTA dopamine neurons for 10-20 min. **D.** Example units in mPFC that are significantly modulated on a prolonged but not transient timescale. Blue rectangles mark the time of stimulation, and * indicates whether the unit was significantly modulated. (See Supplementary Figure 3 for histological confirmation of placements of fiber in VTA and electrode in mPFC and see Supplementary Figure 4 for classification of single units into RS and FS).

We recorded well-isolated single units (*Th::Cre*: n = 131 and wild-type: n = 55) and multi-units (*Th::Cre*: n = 69 and wild-type: n = 24). Most single units recorded in *Th::Cre* rats were classified as regular-spiking (RS) or putative pyramidal units, and only seven units were classified as fast-spiking (FS) (Supplementary Figure 4). The responses of individual units to each phasic VTA stimulation train (either a burst that lasted for 200 ms or a sustained train that lasted for 5 s) on a transient timescale (bin size = 0.05 s for fast and slow burst protocols; bin size = 1 s for sustained phasic protocol) or to the entire stimulation sequence on a longer timescale (bin size = 120 s) was examined. Overall, the response was weak and heterogeneous (Figure 3C,D and 4A,B; Supplementary Table 1,2).

**Figure 4.**
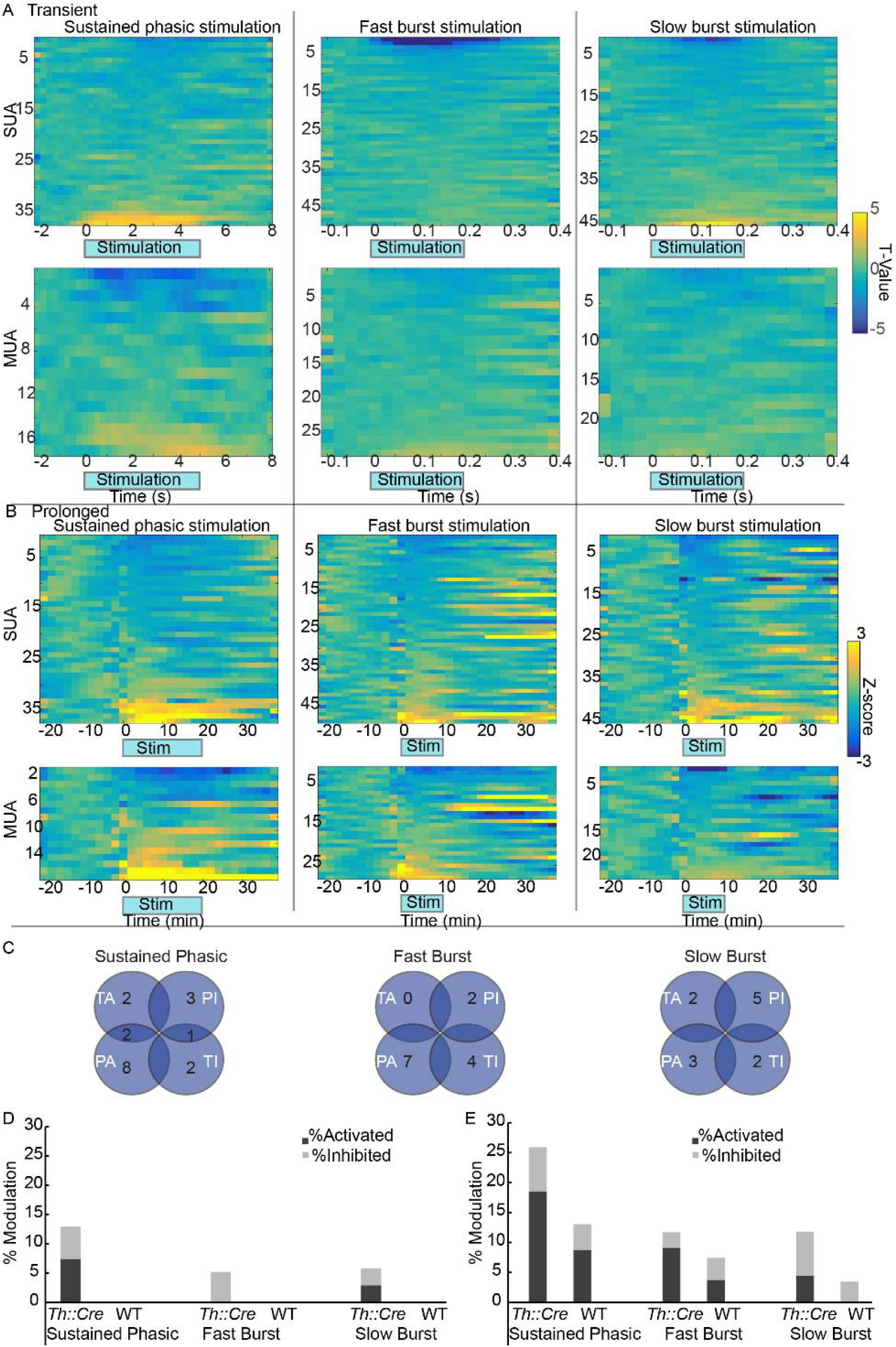
Responses of mPFC units to sustained and burst phasic VTA dopamine activation. **A**. Transient responses of mPFC units. Firing rate changes around each phasic train (that lasted for 5 s in the sustained phasic protocol and for 0.2 s in the fast burst and slow burst protocols) are depicted as t-value color maps for single units (SUA, top) and multi-units (MUA, bottom) recorded in *Th::Cre* rats. Firing rate changes compared to pre-stimulation periods are represented by t-values. Activation and inhibition are indicated by warm and cool colors respectively. Each row indicates one unit, and bin size = 0.025 s for fast and slow burst stimulation and bin size = 0.25 s for sustained phasic stimulation. **B.** Prolonged unit responses to phasic VTA stimulation. Firing rate changes upon stimulation on a prolonged time scale (bin size = 120 s) are depicted as Z-value color maps for single units (SUA, top) and multi-units (MUA, bottom) recorded in *Th::Cre* rats. Firing rate changes compared to the pre-stimulation baseline period are represented by Z-values. Each row indicates one unit. **C**. Venn diagrams summarizing the number of units activated and inhibited on transient and prolonged timescales (TA = transient activated, TI = transient inhibited, PA = prolonged activated, PI = prolonged inhibited) and the overlap in the modulation of activity between those timescales. **D.** Percentage of units (combined SUA and MUA) that were significantly activated/inhibited on a transient timescale for *Th::Cre* and wild-type rats. **E**. Percentage of units (combined SUA and MUA) that were significantly activated/inhibited upon stimulation on a prolonged timescale. For the prolonged timescale analysis, firing rates were analyzed within a window of −10 to 26 min for sustained phasic condition and −10 to −16 min for fast and slow burst conditions.

As shown in specific examples, both excitatory and inhibitory responses in different time scales were observed in RS and FS neurons (Figure 3C,D). For example, in response to sustained phasic stimulation, two example RS units were activated on both transient and prolonged timescales (Figure 3C; transient: top two rows; prolonged: bottom row) whereas two example multi-units were activated or inhibited transiently on the order of seconds but not on the order of minutes. These example responses suggest that mPFC neurons can be modulated either on a transient or a prolonged timescale or both. In fact, the population of units that showed transient versus lasting responses did not overlap much, as only three units across all protocols (and these three units were recorded in the sustained phasic stimulation sessions) showed both significant transient and prolonged responses (Figure 4C), suggesting that different ensembles of neurons may be modulated on different timescales. Thus, modulation of mPFC unit activity over the course of seconds/milliseconds can be independent from modulation over minutes and may arise from different mechanisms.

A summary of modulation across all recorded units is depicted in Figure 4A,B. It is apparent from the color maps that only a handful of units responded transiently and heterogeneously (activation and inhibition) to VTA phasic stimulation (Figure 4A). Overall, only 8% of units (across all stimulation protocols) were transiently modulated by phasic VTA dopamine activity in *Th::Cre* rats. 57% of FS units (compared to 6% of RS units) were modulated by phasic dopamine activity (Supplementary Table 1); the small sample in the present data may, however, render this an unreliable indicator of the effects of dopamine activity on FS units. Nonetheless, it is possible that FS neurons are more likely to be modulated by transient VTA dopamine activity than pyramidal neurons. Combined across all units (single and multi-units), 13% of units were transiently modulated by sustained phasic stimulation, while 5% and 6% of units were modulated by fast and slow burst stimulation respectively (Figure 4D). None of the units (SUA and MUA) recorded in wild-type animals showed significant response to stimulation (Figure 4D and Supplementary Table 1).

On a prolonged timescale, phasic VTA dopamine activity modulated 15% of all recorded units heterogeneously in *Th::Cre* rats (Figure 4B and Supplementary Table 2). Only one of seven recorded FS units was significantly modulated, and this unit’s response is shown in Figure 3D (top panel). This observation, although from a small sample, suggests that maybe FS units are less likely to be modulated on a prolonged rather than a transient timescale. 26% of all mPFC units were modulated by sustained phasic stimulation while 12% and 10% of units were modulated by fast and slow burst stimulation respectively (Figure 4E). Some non-specific unit responses were observed in wild-type rats, possibly reflecting gradual changes in the firing rates of some units over time. A logistic regression model with group and stimulation protocol as predictors (χ^2^ (3) =10.843, p = 0.013) indicated a trend towards difference in modulation between groups (B = 0.808; p = 0.089) and a significant difference in modulation of PFC unit activity by different stimulation types (sustained phasic to fast burst: B = 0.913, p = 0.03; sustained phasic to slow burst; B = 1.162, p = 0.012). Collectively, these data show that the impact of dopamine neuron activation on mPFC individual unit activity is weak, bidirectional (both activation and inhibition), occurs on multiple timescales, and depends on the dopamine neuron activation pattern.

### Modulation of mPFC population response by VTA dopamine activation

Next, we examined activity patterns across the entire mPFC population to better understand how activation of dopamine neurons may modulate neural ensemble response. A notion that has been useful to examine population response is the firing rate or spike count vector ^30, 31^. Each vector consisting of spike counts of N-neurons, which can be denoted by a point in an N-dimensional space, represents the neural population state. The neural population state can change with time and stimulus presentation, and similarity measures, such as Euclidean distances between vectors, can be used to compare population states between different time points or stimulus conditions (Figure 5A).

**Figure 5.**
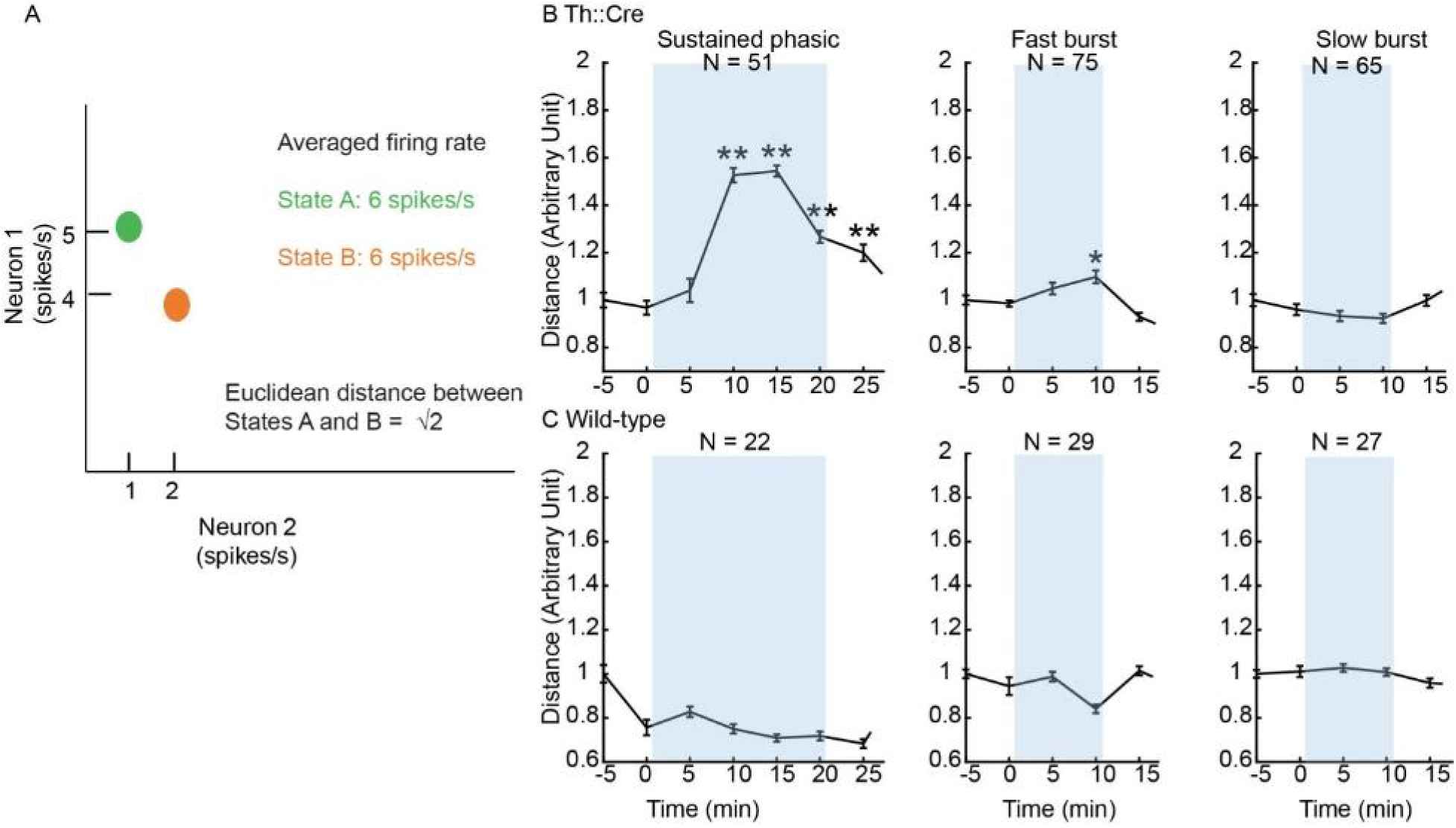
**A**. Illustration of the utility of Euclidean distance measure to assess the similarity/divergence of population states. In this example, the population consists of two neurons, and the spikes/s of these two neurons can be plotted as a point in a two-dimensional space. When stimulus A is presented, Neuron 1 fires 5 spikes/s and Neuron 2 fires 1 spikes/s. When stimulus B is presented, Neuron 1 decreases its firing rate to 4 spikes/s while Neuron 2 increases its firing rate to 2 spikes/s. The changes in firing rate between stimuli A and B are in opposite directions for the two neurons and, therefore, an assessment of population activity by averaging the firing rates of all neurons will indicate that there is no difference in population activity between states A and B (6 spikes/s in states A and B). Further, because the changes in firing rate are small in magnitude (1 spike/s), individual unit responses may not be significantly different between A and B. In contrast, the Euclidean distance measure between states A and B in the 2-dimensional space will accurately indicate that the population responses at A and B are different. This idea can be extended to a population of n-neurons, where each point comprises of spikes/s of n-neurons in the n-dimensional space. **B and C.** Euclidean distances (mean +/− SEM) between population spike count vectors (SUA+MUA, units with firing rate < 0.1 Hz removed) at baseline (time −10 to −15 min before stimulation onset) and all other epochs (each epoch duration = 5 min) are shown for *Th::Cre* and wild-type rats. N indicates the total number of units in each condition, and blue rectangles mark the duration of stimulation (t = 0 corresponds to stim start). To assess the divergence of population activity over time upon stimulation, all distances were compared against the distance between two baseline epochs (−10 to −15 min to −5 to −10 min). Sustained phasic stimulation significantly increased Euclidean distances (compared to baseline) at 5 – 25 min from onset of stimulation sequence (5 – 10 min: p = 0.000; 10 – 15 min: p = 0.000; 15 – 20 min: p = 0.000; 20 – 25 min: p = 0.000; Bonferroni corrected right tailed t-tests). Fast burst stimulation transiently increased Euclidean distance at 5 – 10 min from stimulation onset (p = 0.013), and slow burst stimulation did not modulate Euclidean distances (p > 0.05). Wild-type population activity did not show significant divergence from baseline (all p > 0.05). In the plots, distances are displayed as mean +/− SEM normalized distances (distances for each bin were divided by mean distance between −10 to −15 min and −5 to −10 min epochs). *p < 0.05 and ** p < 0.01.

We utilized the Euclidean distance measure to assess VTA dopamine activity modulation of neural population state over time. For this analysis, we pooled all recorded units to obtain pseudo-spike count population vectors and calculated the spike count vector distance between the baseline period and all other stimulation/post-stimulation periods. We found that upon sustained phasic stimulation of VTA dopamine neurons, mPFC population response exhibited a significant divergence from baseline activity within 5 min from stimulation onset in *Th::Cre* rats but not wild-type rats. This response in *Th::Cre* rats was sustained even at 5 min after stimulation offset, although it was decreased in magnitude by this time point (Figure 5B). Fast burst stimulation of VTA dopamine neurons, on the other hand, produced a transient population response at 5 –10 min from stimulation onset that was significantly different from baseline in *Th::Cre* rats, but the modulation of population response was much smaller in magnitude than with the sustained phasic stimulation (Figure 5B). Slow burst stimulation did not significantly modulate population activity (Figure 5B). Even though fast and slow burst protocols provided the same total duration (= 24 s) of blue laser illumination of VTA in 10 min of stimulation, the 100 Hz versus 20 Hz bursting frequencies elicited different magnitudes of modulation of mPFC population response. Furthermore, even though the sustained phasic stimulation lasted for 20 min (instead of 10 min), the total duration of laser illumination in 10 min (= 30 s) was comparable to that of other protocols; the sustained phasic protocol generated a significant population response within the first 10 min that was much larger than the effect of fast and slow burst stimulation. Thus, phasic activation of dopamine neurons in a sustained pattern elicits a much greater modulation of mPFC population activity than bursting patterns either at the same or higher intra-burst frequencies. These findings suggest that, on a prolonged timescale, repeated phasic activity of VTA dopamine neurons can generate significant population responses in mPFC, and these population-level mPFC neural changes depend on the pattern of dopamine neuron activation.

Phasic VTA dopamine activity did not transiently modulate population-level responses, which were assessed in terms of the similarity of population activity patterns between baseline and stimulation trials (all p > 0.05 for comparison of Euclidean distances between baseline and stimulation trials, Bonferroni corrected; see Methods).

### Behavioral state-dependent modulation of mPFC population response by dopamine activation

Differences in population unit activity (Figure 5) across stimulation protocols were comparable to the differences in the amount of behavioral modulation elicited by those protocols in *Th::Cre* rats (Figure 6). Sustained phasic stimulation significantly and robustly increased movement (movement index tracked all active events, including grooming, rearing, and locomotion) while fast and slow burst stimulation did not change behavior compared to baseline. In wild-type rats, behavior was not significantly modulated by any stimulation protocol (repeated measures ANOVA, sustained phasic stimulation: F(6,18) = 1.372; fast burst stimulation: F(4,12) = 0.640; slow burst stimulation: F(4,4) = 0.805; all p > 0.05).

**Figure 6.**
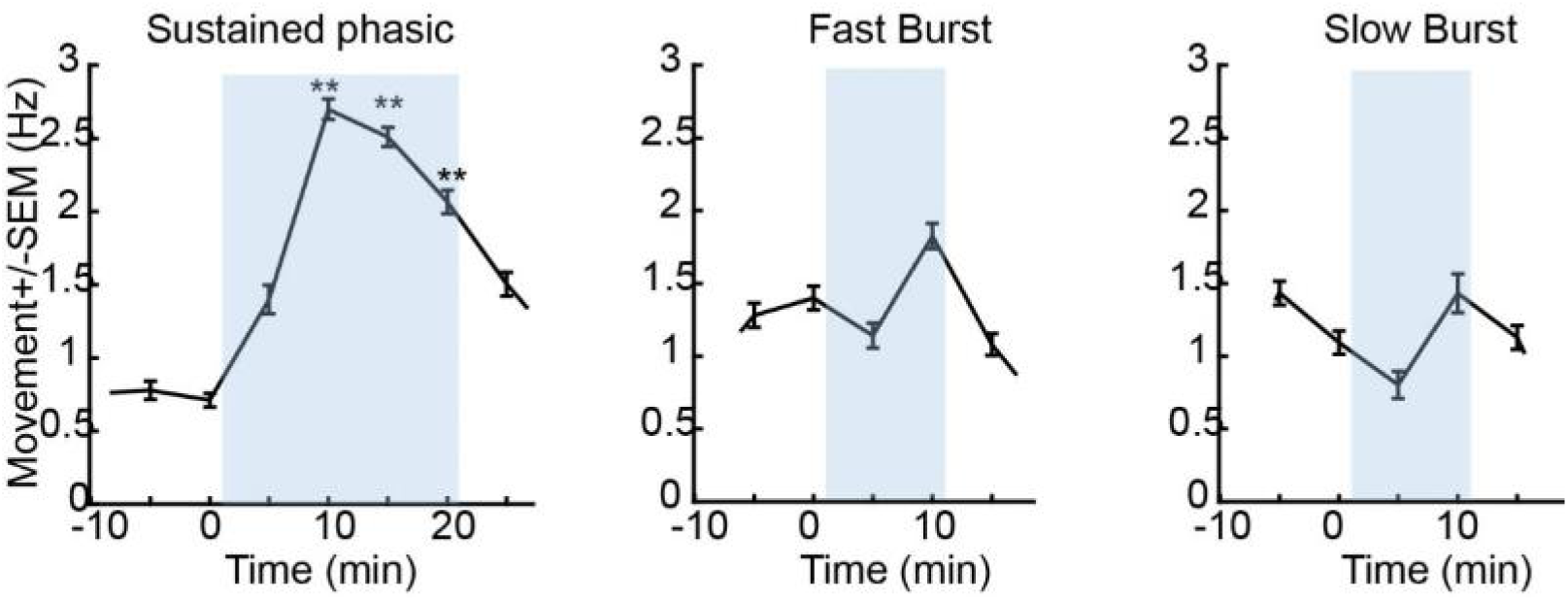
Modulation of spontaneous behavioral activity of *Th::Cre* rats by VTA optogenetic stimulation. Activity or “movement” depicted here includes changes in locomotion, grooming, and rearing as detected with an infrared monitor. Sustained phasic stimulation significantly increased the amount of movement (repeated measures ANOVA: F(6,72) = 8.105, p = 0.000). ** indicates the significance of each bin for comparison against baseline at −10 to −5 min (post-hoc paired t-tests, Bonferroni corrected) at p < 0.01. Fast burst stimulation and slow burst stimulation did not significantly modulate movement (F(4,48) = 1.132, p = 0.184, fast burst; F(4,36) = 0.879, p = 0.474, slow burst). Time = 0 indicates start of stimulation.

Because of the dynamic nature of animal behavior in a freely moving preparation and the observed modulation of behavior by stimulation, we examined PFC population responses during baseline and after stimulation in similar behavioral states. For this, we identified movement epochs in baseline, stimulation, and post-stimulation periods during which animals were moving at comparable rates (Supplementary Figures 5, 6A,6B). Even when the animal’s behavioral state was controlled for, sustained phasic stimulation and fast burst stimulation significantly modulated population responses in the stimulation period in *Th::Cre* rats (Figure 7 & Supplementary Figure 6C). Some of the stimulation-related modulation of population response shown in Figure 5 may be accounted for by an enhancement of animal’s behavioral activity level as the responses in Figure 7 were comparatively weaker. Nonetheless, a significant amount of population activity modulation was independent of potential movement-elicited PFC changes. Accordingly, population response during stimulation occupied a different activity space in PFC compared to movement-elicited PFC response in stimulation-free periods as illustrated in Figure 7 (bottom panels).

**Figure 7.**
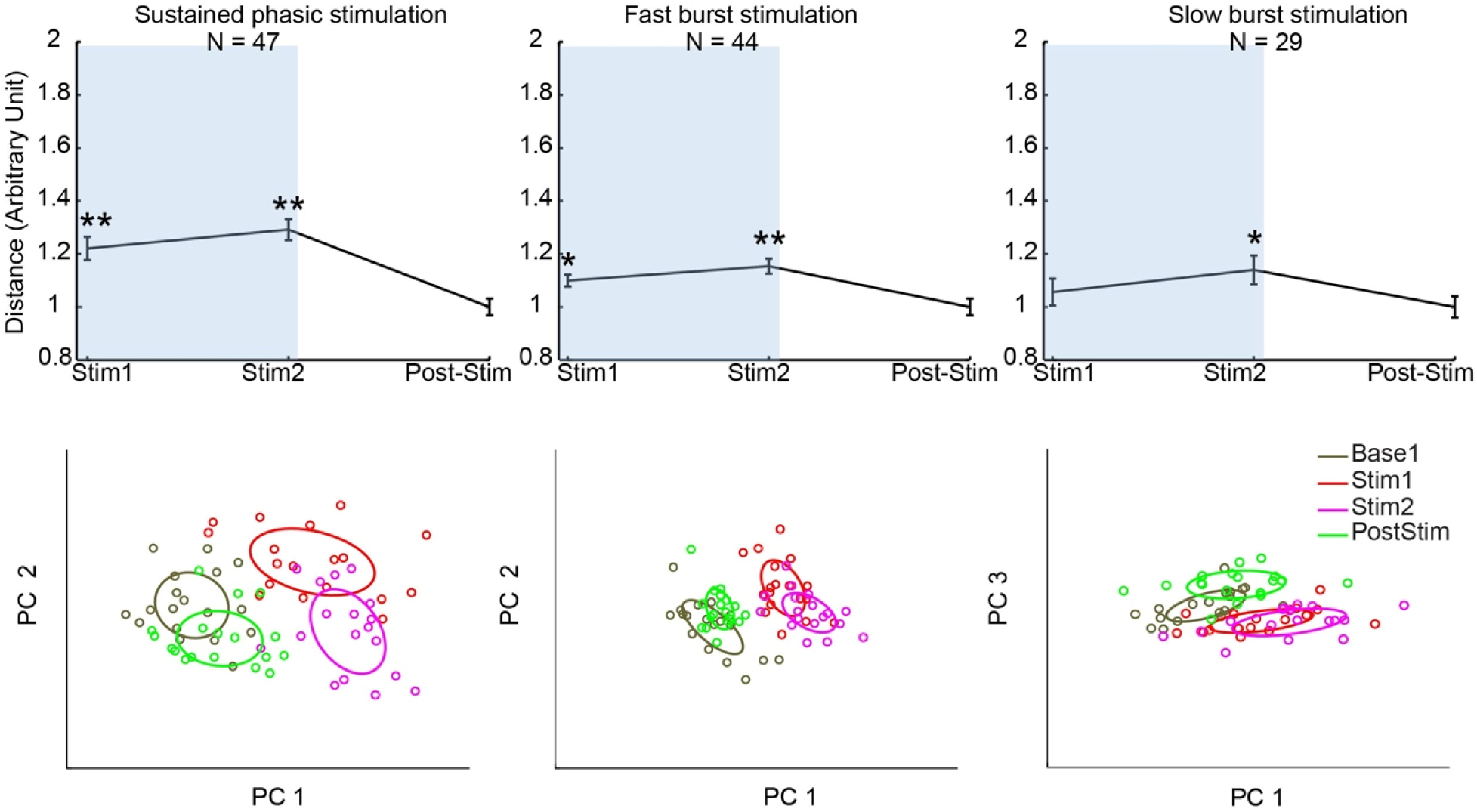
Impact of sustained and burst phasic activation of VTA dopamine neurons on mPFC population activity during behaviorally active epochs in *Th::Cre* rats. Moving epochs (each 3 min long) were identified in baseline, stimulation and post-stimulation periods as shown in Supplementary Figure 5. Stim 1 and 2 correspond to two different moving epochs during the stimulation period, while post-stim indicates an epoch in the period following stimulation termination. Euclidean distances of population spike count vectors (SUA+MUA, and units with firing rate < 0.1 Hz removed) between baseline and all other epochs are shown for all stimulation protocols. To assess the divergence of population activity over time upon stimulation, all distances were compared against the distance between baseline and post-stim epoch. Sustained phasic stimulation significantly increased population activity divergence from baseline during stimulation epochs (Stim 1: p = 0.000; Stim 2: p = 0.000; Bonferroni corrected one tailed t-tests). Fast burst stimulation increased population divergence in all stimulation epochs (Stim1: p = 0.015, Stim2: p = 0.000), and slow burst stimulation only weakly modulated Euclidean distance at Stim 2 (p = 0.046). To allow visualization of this divergence of population activity during stimulation, in bottom panels, high dimensional spike count vectors were analyzed with PCA and projected onto the space of any two of the first three principal components that resulted in maximal separation between baseline and stimulation epochs. Ellipses indicate covariance within each epoch. In the distance plots, distances are displayed as mean +/− SEM normalized distances (distances for each bin were divided by the mean distance between baseline and post-stimulation epochs). *p < 0.05 and ** p < 0.01. N indicates the total number of units in each condition, and blue rectangles mark stimulation epochs.

While significant population responses to slow burst stimulation were not detected in the analysis in Figure 5, there was a weak modulation of population activity in one of the moving stimulation epochs (Figure 7). This discrepancy may indicate that dopamine modulation of population response in PFC is state dependent and is more likely to occur if the animal is behaviorally active.

### Modulation of mPFC local field potentials (LFPs) by phasic VTA dopamine activity

Next, we examined how phasic activation of VTA dopamine neurons modulated LFPs in mPFC. LFP frequencies were binned into several bands: delta (1 – 4 Hz), low theta (4.5 – 8 Hz), high theta/alpha (8 – 13 Hz), beta (14 – 30 Hz), low gamma (30 – 55 Hz), and high gamma (55 – 100 Hz), and power modulation in these bands was assessed.

On a transient timescale, VTA stimulation with sustained phasic and slow burst paradigms significantly enhanced PFC beta power in *Th::Cre* rats (Figure 8). Fast burst stimulation did not modulate beta power. In wild-type animals, none of the stimulation paradigms elicited a significant modulation of beta power (Sustained phasic: F(9,27) = 0.638, p = 0.755; Fast burst: F(9,27) = 2.214, p = 0.053; Slow burst: F(9,18) =0.958, p = 0.490; repeated measures ANOVA) (Supplementary Figure 7). Because of the low sample size in each wild-type stimulation group, we also pooled data from sustained phasic and slow burst sessions (both stimulation protocols use 20 Hz frequency) and analyzed changes in power within −0.1 to 0.4s of each stimulation train. The pooled data did not show significant beta power modulation by 20 Hz light stimulation alone (F(9,54) = 1.360, p = 0.257) (Supplementary Figure 7D). In contrast, when data from sustained phasic stimulation sessions of *Th::Cre* rats were analyzed with the same window (−0.1 to 0.4s), significant modulation of beta power was still observed (F(9,126), p = 0.002). To rule out the possibility that the effects observed may be an artifact of stimulation frequency, we examined LFP power changes up to 120 Hz during fast burst stimulation (that utilizes 100 Hz stimulation frequency) and did not observe any modulation in frequencies around 100 Hz (Supplementary Figure 7E). Thus, VTA stimulation with sustained phasic and slow burst paradigms transiently and selectively enhanced beta activity in PFC LFPs.

**Figure 8.**
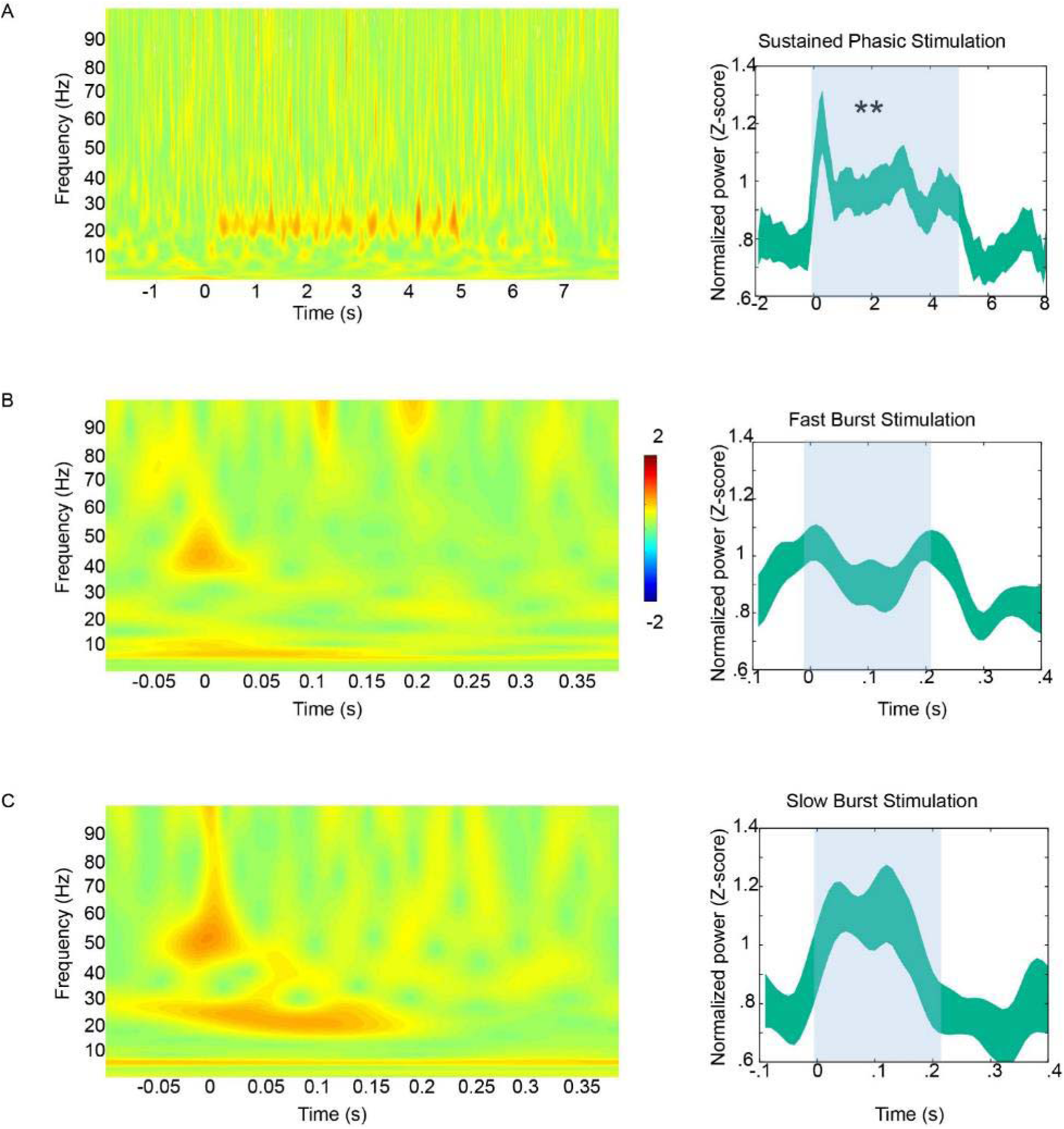
Transient modulation of LFPs by phasic VTA dopamine activity in *Th::Cre* rats. (Left panels) LFP spectrograms (normalized to total LFP power within each frequency bin) depict responses to each phasic train in in the corresponding stimulation protocol. Color bar indicates Z-scores. (Right panels) Mean +/− SEM normalized beta frequency power in different time bins. T = 0 corresponds to stimulation train onset and blue rectangles mark the duration of each stimulation train. **A.** Sustained phasic stimulation significantly increased beta (14-30 Hz) power (F(9,126) = 10.781, p = 0.000; repeated measures ANOVA), and post-hoc comparisons indicated significant elevations in power at time t = 0 – 1 s (p = 0.028, Bonferroni corrected paired t-test), and t – 2-3 s (p = 0.008). Transient modulation in other frequency bands was not observed (p > 0.05, repeated measures ANOVA). **B.** Fast burst stimulation did not modulate beta band power (F(9,126) = 1.570, p = 0.167; repeated measures ANOVA) or any frequency band transiently (p > 0.05 for all frequencies). **C.** Slow burst stimulation increased beta frequency band, and the increase showed a trend toward significance (F(9,99) = 2.451, p = 0.080, repeated measures ANOVA). Other frequency bands were not affected (p > 0.05). * and ** indicate p < 0.05 and p < 0.01 respectively for repeated measures ANOVA.

VTA dopamine stimulation resulted in a different pattern of PFC LFP modulation on a prolonged timescale (Figure 9). The spectrograms in Figure 9 (top panels) depict power changes over the course of the entire session. Statistical analysis (Figure 9, bottom panels) was restricted to periods closer to the stimulation epoch to ensure a similar behavioral state, characterized by mostly awake, alert epochs, across time points. As is apparent in the spectrograms, the baseline period prior to −10 min and the post-stimulation period beyond 20 min (from stimulation onset) in all three paradigms was characterized by long periods of enhanced lower frequency activity and reduced high frequency oscillations, which is indicative of a drowsy or sleepy rat ^32^.

**Figure 9.**
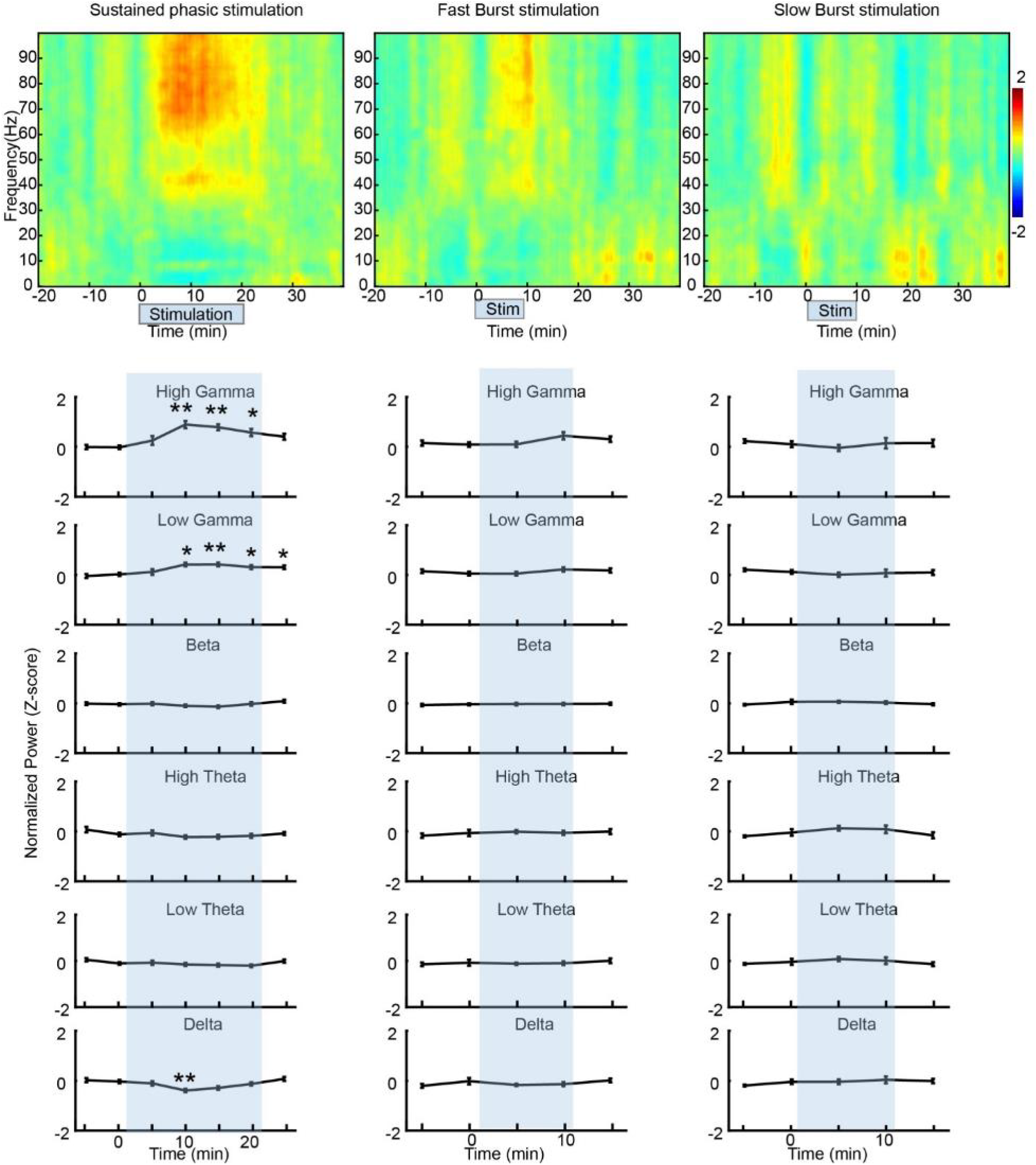
Phasic activation of VTA dopamine neurons modulated LFPs on a prolonged timescale in *Th::Cre* rats. (Top panels) Power spectrograms depict Z-score normalized LFP power for frequencies between 0 – 100 Hz before and after stimulation with sustained phasic, fast burst, and slow burst protocols. Color bar indicates Z-scores. (Bottom panels) Mean +/− SEM normalized power values for different frequency bands at different time bins (bin width = 5 min). T = 0 min corresponds to stimulation onset and blue rectangles mark the duration of stimulation. Sustained phasic stimulation significantly increased high gamma (55 – 100 Hz) power (F(6,84) = 13.08, p = 0.00) at 5-10 min (p = 0.002; Bonferroni corrected t-test), 10-15 min (p = 0.000), and 15-20 min (p = 0.036) from the onset of stimulation. Sustained phasic stimulation also significantly increased low gamma (30 – 55 Hz) power (F(6, 84) = 7.019, p = 0.000) at 5-10 min (p = 0.011), 10-15 min (p = 0.008), 15-20 min (p = 0.038), and 20-25 min (p = 0.028) and decreased delta (1 – 4 Hz) power at 5-10 min (p = 0.009) from stimulation onset. Beta (F(6,84) = 2.248, p = 0.060), high theta (F(6,84) = 1.896, p = 0.111), and low theta (F(6,84) = 2.233, p = 0.072) power were not modulated by sustained phasic stimulation. Fast burst stimulation elicited a trend towards significant increase in high gamma power (F(4,56) = 2.176, p = 0.096). Low gamma (F(4,56) = 0.914, p = 0.453), beta (F(4,56) = 0.204, p = 0.922), high theta (F(4,56) = 0.529, p = 0.699), low theta (F(4,56) = 0.411, p = 0.754), and delta (F(4,56) = 1.111, p = 0.250) frequencies were not modulated by fast burst stimulation. Slow burst stimulation did not significantly modulate power in any frequency band (high gamma: F(4,44) = 0.620, p = 0.448; low gamma: F(4,44) = 0.656, p = 0.536; beta: F(4,44) = 0.864, p = 0.476; high theta: F(4,44) = 1.512, p = 0.231; low theta: F(4,44) = 0.748, p = 0.497; delta: F(4,44) = 0.751, p = 0.513). * and ** indicate p < 0.05 and p < 0.01 for comparison of each bin against baseline time point −10 to −5 min (post-hoc paired t-tests; Bonferroni corrected).

Upon sustained phasic VTA dopamine activation, the most prominent increase in LFP power was detected in the broadband high gamma (55 – 100 Hz) frequency band (Figure 9). High gamma power increased within 10 min of stimulation onset and persisted until the end of stimulation (mean Z-scored power at significant samples = 5-10 min (0.896 +/− 0.144), 10 - 15 min (0.7872 +/− 0.118), and 15 - 20 min (0.5712 +/−0.145)). Sustained phasic stimulation also increased power in the low gamma (30 – 55 Hz) frequency band, although the increases were weaker in magnitude (mean Z-scored power at significant samples = 5 - 10 min (0.424 +/− 0.075), 10 - 15 min (0.427 +/− 0.079), 15-20 min (0.318 +/− 0.079), and 20 - 25 min (0.314 +/− 0.071)) compared to the high gamma band. Fast burst stimulation only resulted in a trend towards significant modulation of high gamma power, and this trend may have been driven by a small increase in high gamma at 5 - 10 min (Z-score = 0.443 +/− 0.146); other frequency bands were not significantly modulated. Slow burst stimulation failed to elicit significant changes in any frequency band.

Despite modulation of gamma power, our analyses did not reveal a significant enhancement of phase-locking of mPFC neuronal spikes to local gamma oscillations. While there was some increase in the proportion of units phase-locked to high gamma frequency during sustained phasic stimulation (Supplementary Figure 8), the differences in proportions were not significant (p > 0.05; Fisher’s exact test). There was no difference in the proportion of units phase-locked to low gamma frequency band (p > 0.05; all Fisher’s exact tests). There was also no difference in phase locking value (PLV), which measures the strength of phase-locking, for all units before and after stimulation for gamma bands (Friedman’s ANOVA, p > 0.05). Similarly, stimulation with fast and slow burst protocols did not elicit a significant change in the proportion of phase locked units (p > 0.05; Fisher’s exact tests) and PLVs for gamma frequencies (p > 0.05; Friedman’s ANOVA). The absence of a significant phase locking relationship may be due to our electrodes not recording from enough units that were phase-locked to the oscillatory activity or that dopamine-induced oscillatory activity in PFC is caused by mechanisms other than what is assumed conventionally^33^.

Modulation of high gamma power by sustained phasic stimulation persisted even when the behavioral state was controlled for by analyzing data from moving epochs only (Figure 10). This modulation, however, was weaker than when data from the entire session was analyzed (Figure 9). Similar to the effects on population activity, fast and slow burst stimulation resulted in a greater modulation of high gamma power when animals were behaviorally active (Figure 10) compared to when animal behavior was not factored in (Figure 9). These results demonstrate that VTA dopamine activation modulates high gamma power, and this modulation of power shows an interaction between the pattern of dopamine activation and the behavioral state of the animal.

**Figure 10.**
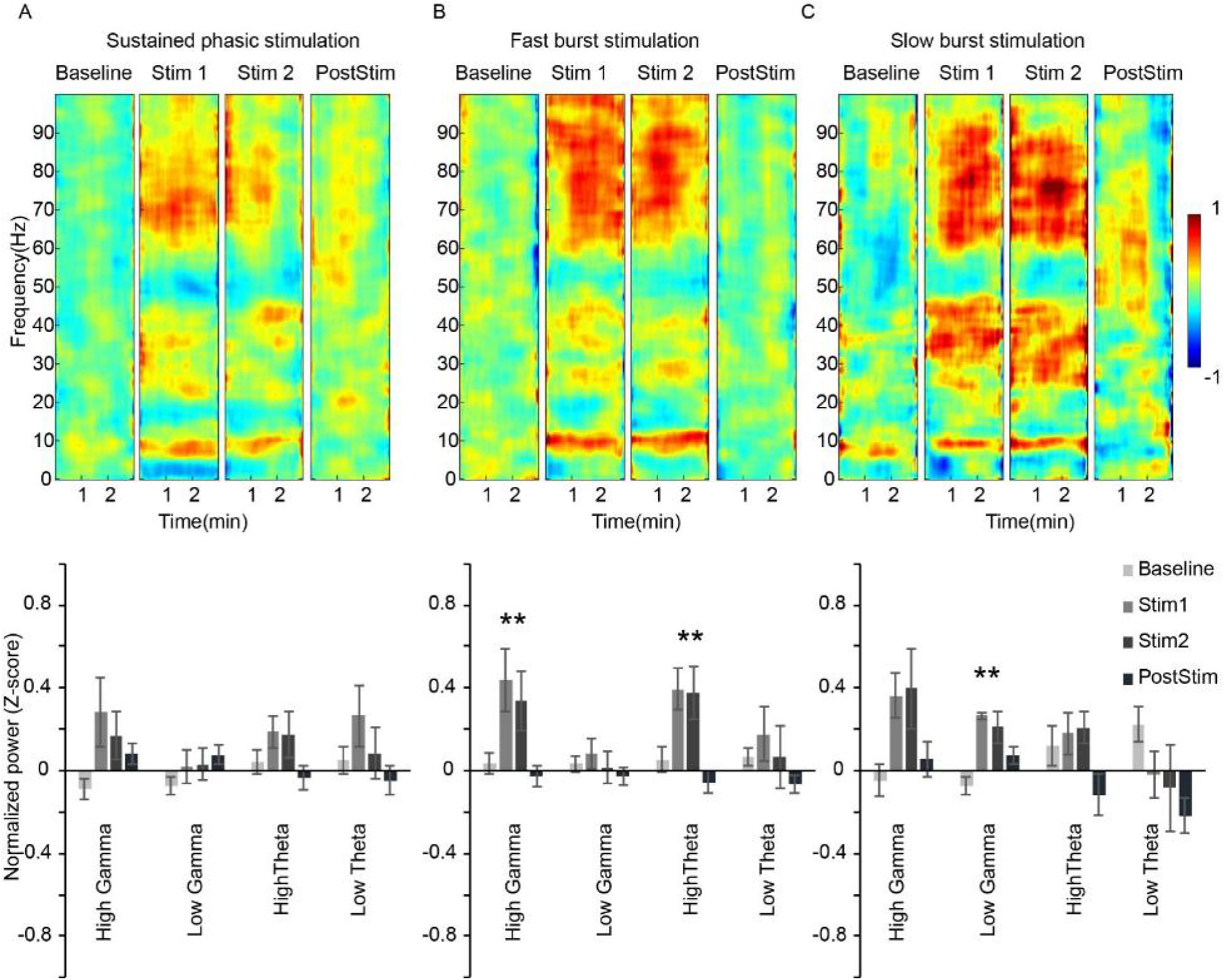
Impact of sustained and burst phasic activation of VTA dopamine neurons on mPFC LFPs during behaviorally active epochs in *Th::Cre* rats. Moving epochs (each 3 min long) were identified in baseline, stimulation and post-stimulation periods as shown in Supplementary Figure 5. Stim 1 and 2 correspond to two different moving epochs during the stimulation period. Power spectrograms (top panels) depict LFP power for frequencies within each epoch as Z-score normalized values. Color bar indicates Z-scores. Bar plots (bottom panels) show mean +/− SEM of normalized Z-score power values for high gamma, low gamma, high theta, and low theta frequency bands. **A** Sustained phasic stimulation increased the power of high gamma with a trend towards significance (F(3,30) =2.661, p = 0.066; repeated measures ANOVA). **B** Fast burst stimulation significantly increased high gamma power (F(3,24)=4.890, p = 0.009) and high theta power (F(3,24) = 6.534, p = 0.002). Post-hoc tests revealed a significant increase in high gamma in stim epoch 1 (p = 0.012), and a trend towards significant increase in stim epoch 2 (p = 0.086). High theta power was elevated in stim epoch 1 (p = 0.038) and stim epoch 2 (p = 0.055). **C** Slow burst stimulation significantly increased low gamma power (F(3,9) = 8.283, p = 0.006) and resulted in a trend towards significant increase in high gamma power (F(3,9)= 3.656, p =0.057). Post-hoc tests revealed a significant increase in low gamma power in stim epoch 1 (p = 0.002) and a trend in stim epoch 2 (p = 0.061). * and ** indicate significance of repeated measures ANOVA at p < 0.05 and p < 0.01 respectively.

VTA stimulation did not modulate LFP power in wild-type rats in the entire session analysis (sustained phasic stimulation: F(6,18) = 0.960 (high gamma), F(6,18) = 0.694 (low gamma), F(6,18) = 0.895 (beta), F(6,18) = 0.668 (high theta), F(6,18) = 0.772 (low theta), and F(6,18) = 0.715 (delta); fast burst stimulation: F(4,12) = 0.814 (high gamma), F(4,12) = 0.794 (low gamma), F(4,12) = 0.339 (beta), F(4,12) = 0.517 (high theta), F(4,12) = 0.699 (low theta), F(4,12) = 0.610 (delta); slow burst stimulation: F(4,8) = 1.455 (high gamma), F(4,8) = 2.752 (low gamma), F(4,8) = 1.367 (beta), F(4,8) = 1.886 (high theta), F(4,8) = 2.225 (low theta), and F(4,8) = 2.471 (delta); all p > 0.05). Modulation of LFP power was also not observed when data from moving epochs only were analyzed (Supplementary Figure 9).

### Modulation of phase amplitude coupling by phasic dopamine neuron activity

Next, we examined if and how the phase of slower PFC LFP frequencies (delta, low theta or high theta) modulated the amplitude of higher LFP frequencies (low gamma or high gamma) upon dopamine neuron activation. An example of how the phase of theta oscillations can modulate the amplitude of gamma oscillations is illustrated in Figure 11A. Sustained phasic VTA stimulation, in addition to increasing PFC high gamma power, also increased the coupling between PFC high gamma and theta oscillations (Figure 11B), in both high and low theta ranges (high theta to high gamma coupling: χ^2^(6) = 14.029, p = 0.029, Friedman’s ANOVA; post-hoc: n.s., signed-rank tests, Bonferroni corrected; low theta to high gamma: χ^2^(6) = 24.91, p = 0.000, Friedman’s ANOVA; post-hoc: p (5 – 10 min) = 0.003; p (10 – 15 min) = 0.062, signed rank with Bonferroni correction). These effects were not spurious changes due to random coupling by chance or simply because of increased power because surrogate LFPs (n = 100 surrogate time series obtained by splicing the time series into two random splits within each bin) did not show such coupling (low theta to high gamma: χ^2^ (6) = 5.714, p = 0.456; high theta to high gamma: χ^2^ (6) = 3.686, p = 0.719). Similarly, there was a significant increase in the coupling between high theta and high gamma upon fast burst stimulation (χ^2^ (4) = 9.867, p = 0.043, post-hoc: n.s., signed-rank tests, Bonferroni corrected; surrogate analysis: χ^2^ (4) = 1.547, p = 0.818). Slow burst stimulation did not modulate coupling between any of the frequencies tested (all p > 0.05 for Friedman’s ANOVA). Furthermore, there was no change in the coupling of delta and high gamma or delta and low gamma frequencies after VTA dopaminergic stimulation with any protocol (all p > 0.05 for Friedman’s ANOVA), in contrast to the observation of a previous study ^34^. Therefore, during epochs when significant increases in high gamma power were observed upon sustained phasic stimulation and a trend towards increase was observed upon fast burst stimulation, there was also a corresponding increase in the coupling between theta and high gamma frequencies. There was no significant phase-amplitude coupling between any frequencies tested for wild-type animals after sustained phasic stimulation (n = 4 rats), fast burst stimulation (n = 4 rats), or slow burst stimulation (n = 3 rats) (all p > 0.05, Friedman’s ANOVA).

**Figure 11.**
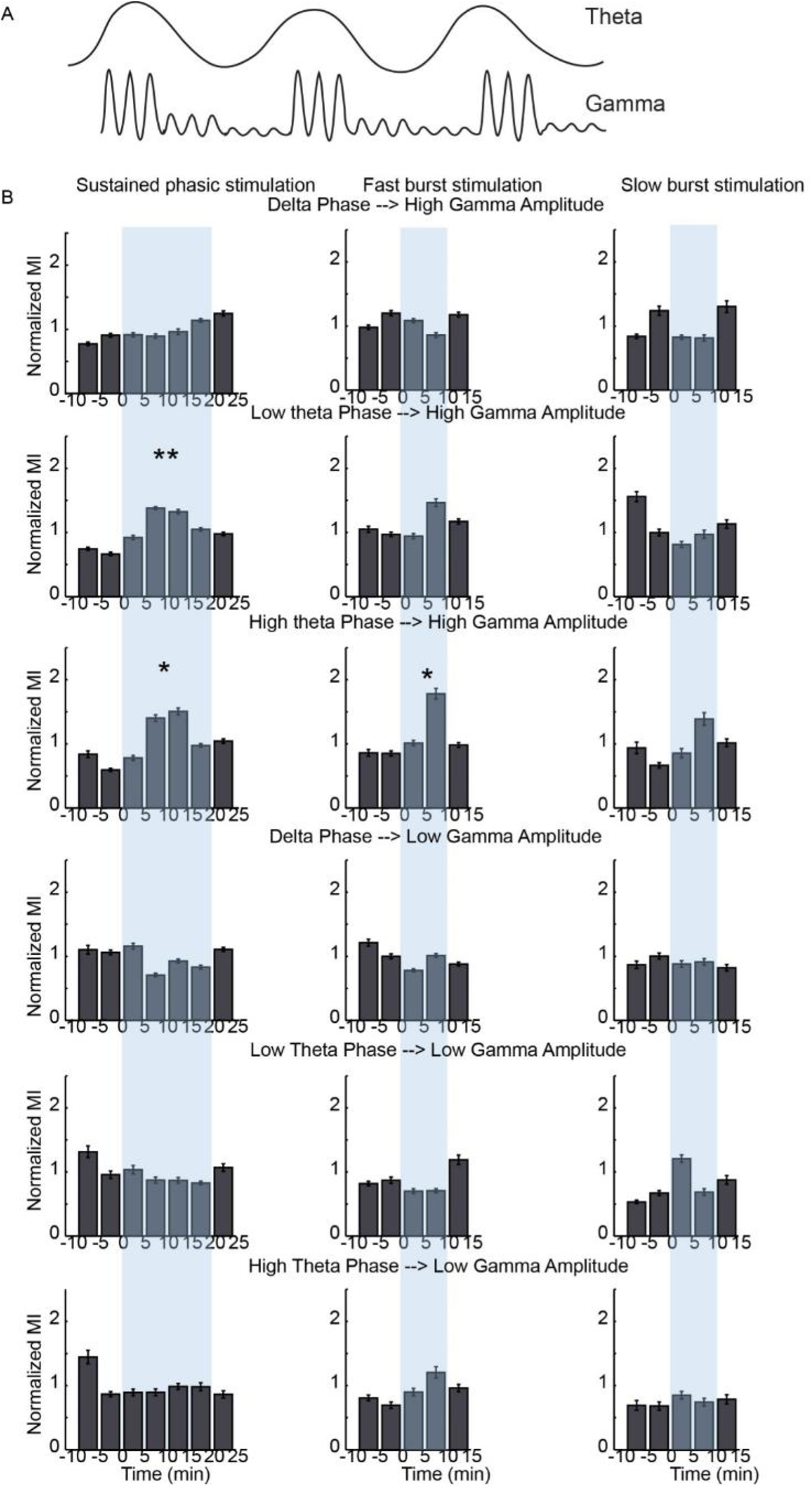
**A**. Schematic of phase amplitude coupling. Illustrations of filtered LFP traces at theta and gamma ranges are shown to depict how the phase of a low frequency oscillation can modulate the amplitude of a high frequency oscillation. **B.** Mean +/−SEM normalized modulation index (MI) values of phase-amplitude coupling in *Th::Cre* rats (sustained phasic: n = 15 rats, fast burst: n = 15 rats, slow burst: n = 12 rats). MI values are plotted across time to depict phase-amplitude coupling between delta and high gamma, low theta and high gamma, high theta and high gamma, delta and low gamma, low theta and low gamma, and high theta and low gamma for *Th::Cre* rats. T = 0 indicates stimulation onset, * and ** indicate significance of Friedman’s ANOVA at p < 0.05 and p< 0.01 respectively.

Theta-high gamma coupling modulation persisted when movement epochs were selectively analyzed. Enhanced coupling between high theta and high gamma during fast burst stimulation (χ^2^(3) = 17.667, p = 0.000; post-hoc: p (Stim 1) = 0.020, p (Stim 2) = 0.004; surrogate analysis: χ^2^(3) = 0.333, p = 0.953) and a trend toward significant increase in coupling between high theta and high gamma during slow burst stimulation (χ^2^ (3) = 6.600, p = 0.086; surrogate analysis: χ^2^(3) = 4.500, p = 0.212) were observed (Figure 12). Similar to the effect on gamma power of LFPs (Figure 10), the impact of slow burst stimulation on theta-gamma cross frequency coupling was selectively unmasked during analysis of behaviorally active epochs (Figure 12) compared to when behavior was not accounted for (Figure 11). No modulation of cross frequency coupling was observed in wild-type animals (all p > 0.05, Friedman’s ANOVA; n = 6 sessions, data were pooled across protocols). These findings suggest that VTA dopamine activation increases the coupling between theta and high gamma frequencies. This effect is associated with high gamma power elevation and depends on the pattern of dopamine neuron activation and animal behavioral state.

**Figure 12.**
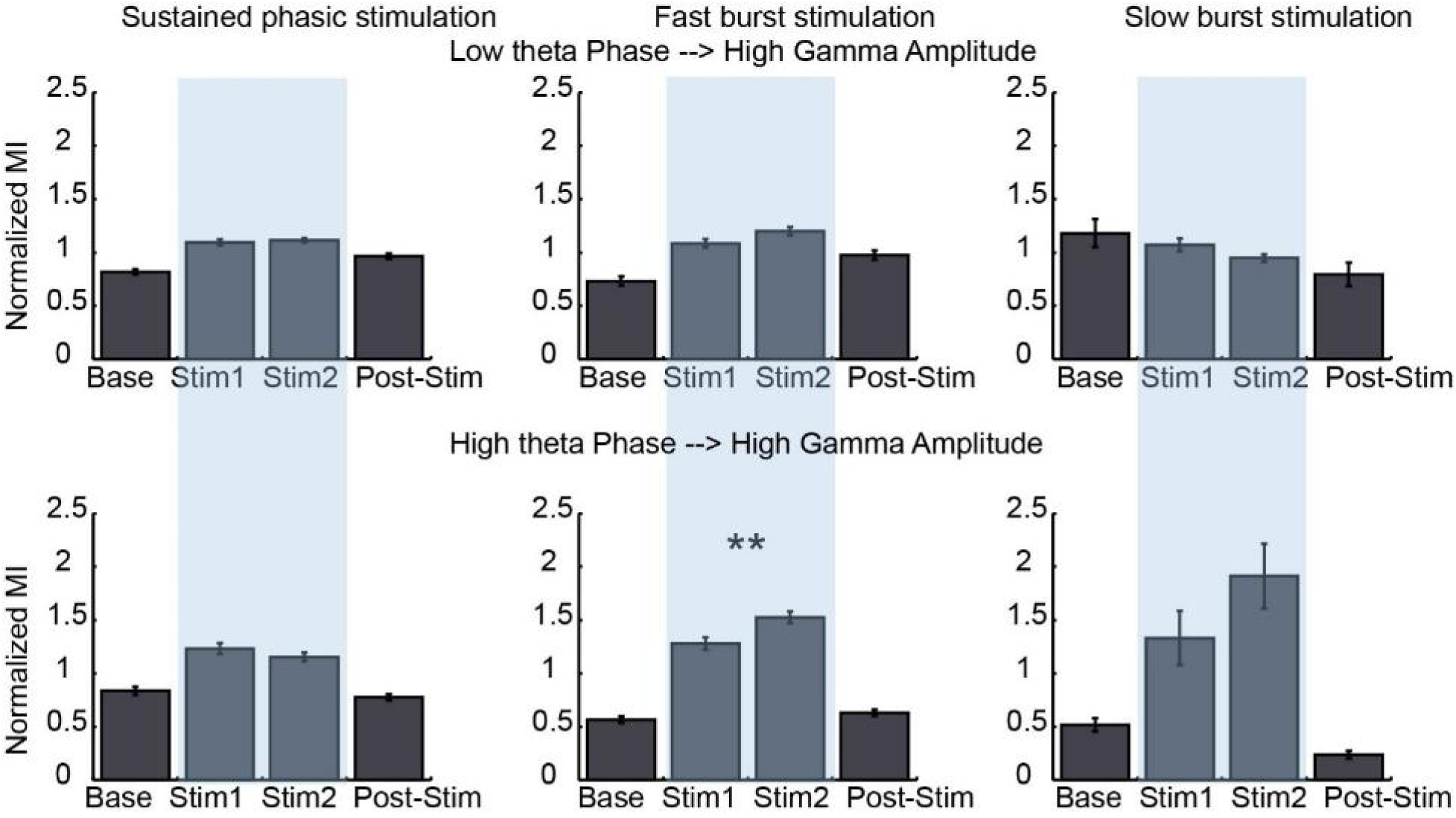
Mean +/−SEM normalized modulation index (MI) values for behaviorally active epochs in *Th::Cre* rats (sustained phasic: n = 12 rats, fast burst: n = 9 rats, slow burst: n = 4 rats). Moving epochs were identified as described in Supplementary Fig 5. * and ** indicate significance of Friedman’s ANOVA at p < 0.05 and p < 0.01 respectively.

## Discussion

We wanted to understand how dopamine can modulate PFC computation of temporally diverse events. Activation patterns of dopamine neurons are not uniform and are highly sensitive to context ^35^. While a classic tonic versus phasic pattern of firing has long been reported in ex-vivo and anesthetized preparations ^36-38^, in behaviorally relevant contexts, the phasic response patterns of these neurons are complex and heterogeneous ^23, 26, 39^. As typified by the raw trace examples shown in Figure 1, dopamine neurons display both sustained ^23, 40^ and burst ^22, 40, 41^ patterns of phasic responses during active behavior. In addition, while some of dopamine related functions may operate on a trial by trial basis, task performance requires sustained motivation and behavioral engagement/attention ^42^ for many minutes that may be maintained by PFC computations dependent on prolonged phasic VTA dopamine activity. To model these circumstances, we used optogenetics in freely moving rats to stimulate VTA dopamine in different patterns over a typical period (10-20 minute) of task performance as we measured spontaneous unit activity and LFPs in PFC along with behavior. We found that the impact of dopamine on individual unit discharge rate was weak and heterogeneous in directionality and duration. In contrast, we observed a robust and prolonged impact on coordinated ensemble activity, gamma oscillations and gamma-theta coupling. Sustained phasic activation of dopamine neurons, which has been reported during tasks involving sustained attention and distant-goals ^23, 24^ and is distinct from the classical and frequently reported burst responses, consistently modulated PFC ensemble and LFP activity. The phasic burst pattern of dopamine activation, which is typically associated with salience detection and reward prediction error coding ^22, 40, 41^, had the most profound impact on PFC network activity during active behavior. Collectively, these results demonstrate causation between phasic activity of dopamine neurons and multiple spatio-temporal modes of modulation of PFC activity that occur concomitantly. These data further provide evidence that dopamine may support diverse computations by PFC at the service of multiple and temporally varied functions such as working memory, reward-guided learning, action selection, motivation, anxiety states, behavioral inhibition, and attentional tuning.

### >Impact of sustained or burst activation of dopamine neurons on PFC spontaneous spike activity

The overall effect of dopamine cell firing on spontaneous PFC unit activity was weak and heterogeneous both in terms of the duration and the direction of response. A small percentage of units showed either inhibition or excitation in transient and/or prolonged timescales. These observations are consistent with dopamine’s role as neuromodulator and reports of both excitatory and inhibitory effects of dopamine on PFC neurons ^20, 42-45^. Despite the slow nature of dopamine clearance and extrasynaptic localization of dopamine receptors in PFC ^15, 16^, we observed that some PFC neurons responded on the order of milliseconds to seconds to VTA dopamine neuron stimulation. This fast response is consistent with potential co-release of glutamate in a subset set of these neurons^42, 46, 47^. Finally, we observed that the same PFC cell type (FS or pyramidal) can respond to dopamine stimulus in different ways. This is a novel and unexpected observation that is contrary to a prevalent theory in the field that VTA dopamine activity primarily enhances the firing of FS interneurons and indirectly inhibits the firing of pyramidal neurons ^20^. Our findings, thus, demonstrate that phasic dopamine cell firing modulates the firing rate of some individual PFC units on the timescale of milliseconds to many minutes. This wide time span of response may be relevant to the temporal complexities of several roles that are assigned to PFC dopamine.

### VTA dopamine neuron activation has a robust influence on PFC ensemble activity

Assessing the computational capacity of ensembles of neurons can be difficult to decipher from considering the rates of individual neurons alone. While classic averaging of population responses of groups of neurons can provide useful data in some contexts, the weak and mixed responses (in direction and duration) of individual PFC unit activity to dopamine neuron stimulation made that approach less meaningful for the present data. Thus, we used a population coding approach ^30, 31, 48^ that uses spike count vector to compare population states between different time points or stimulus conditions. This geometric approach can reveal hidden patterns of neural ensemble activation and has been used successfully in sensory systems; for example, in the olfactory system, this method has demonstrated that the post-synaptic response to a given stimulus is not linearly scaled but is progressively differentiated in time ^31^. The present study specifically used a Euclidean distance measure to compare the population neural state at different time points/conditions. We observed that the impact of dopamine neuron activation on modulation of PFC activity was best captured by this computation in that sustained phasic and fast burst activation significantly increased Euclidean distance starting at 5 minutes from stimulation onset. This suggests that while activation of dopamine neurons may have a rapid and precise impact on a few PFC units, the most prominent influence on PFC neural ensembles emerges progressively later. The effect of sustained phasic activation on population activity persisted until 5 min after stimulation offset. This lasting mode of modulation of PFC population activity patterns by dopamine may facilitate and strengthen information processing in PFC over many minutes.

Dopamine’s impact on PFC population activity was influenced by the behavioral state of the animal. Initially, we observed that specifically sustained mode, rather than burst mode, of phasic dopamine activation produced a pronounced increase in PFC population response. Subsequent comparison of PFC activity between baseline and stimulation epochs with comparable levels of active behavior yielded a significant modulation of this ensemble response by burst modes of dopamine firing as well. This suggests that prolonged PFC population response to phasic dopamine firing is sensitive to the behavioral state of the animal and may in some contexts emerge more robustly when animals are behaviorally active. These data provide a computational mechanism for how specific patterns of dopamine cell firing, depending on the behavioral context, can contribute to seconds-minutes long cognitive processes such as sustained attention and working memory, and modulation of motivational and affective states.

### VTA dopaminergic modulation of mPFC oscillations

We found a causative relationship between phasic VTA dopamine activity and enhancement of PFC gamma oscillations, more robustly at high gamma range compared to low gamma in the timescale of minutes. Gamma response in PFC to dopamine activation was affected by the behavioral state of the animal in that burst modes of dopamine firing strongly enhanced high gamma selectively in behaviorally active states. Both high and low gamma oscillations are implicated in dopamine-dependent cognitive processes such as working memory and attention ^49, 50^. High gamma is emerging as the most impaired oscillatory range in patients of psychiatric disorders especially schizophrenia ^51^ and relevant genetic models of the disorder ^52^. Thus, our finding that phasic activation of dopamine neurons in an awake preparation is sufficient to enhance high gamma oscillations may have important implications for the pathophysiology of these disorders.

A hypothesized function of LFP oscillations is to synchronize spikes so that they arrive in postsynaptic areas in coherent time windows and enable effective communication ^53^. We, however, found that an increase in high gamma power was not associated with an enhanced gamma-spike entrainment. A similar dissociation has been previously observed during dopamine-dependent hyperkinesia states ^54^. Dopamine modulation of high gamma oscillations, in the absence of local spike synchronization, may tune mPFC to receive afferent inputs in specific time windows ^53, 55, 56^ rather than to modulate the temporal dynamics of outgoing spikes. This can serve as a filtering mechanism and enhance signal to noise.

In addition to facilitating inter-areal communication in specific frequencies, LFP oscillations can interact via cross frequency coupling ^57^, which may be important for dopamine-dependent cognitive functions such as working memory and learning ^50, 58-60^. We found that phasic VTA dopamine activity enhanced high gamma coupling to mPFC theta oscillations during periods when high gamma power was observed to be elevated. This suggests two possibilities: a) that activation of VTA dopamine neurons could enhance the amplitude of PFC high gamma oscillations by modulating the mechanism mediating coupling between theta and high gamma, or b) that dopamine activation could provide multiplexed code to mPFC by independently modulating high gamma power and organizing gamma cycles around theta. The latter form of multiplexing is thought to be important in spatial working memory ^60^ and may explain the nature of the longstanding and necessary role for cortical dopamine in this cognitive construct ^1^.

On a transient timescale, dopamine activation generated a different pattern of LFPs in that sustained phasic and slow burst patterns but not fast burst enhanced PFC beta activity. These beta changes were time-locked to the duration of each phasic train and did not persist for minutes. Because of the selective enhancement of this frequency band by dopamine stimulation in 20 Hz but not 100 Hz frequencies, we were concerned that this may be an artifact of stimulation. This effect, however, was not observed in wild-type animals. Furthermore, the recording electrode was placed in PFC, which is very distal from VTA where light stimulation occurred. The differential enhancement of beta versus gamma band in transient and prolonged timescales may be due to the involvement of different postsynaptic dopamine receptors. Further studies are needed to test the involvement of different mechanisms in different timescales of dopamine activation.

### Behavioral state modulates the impact of dopamine neuron burst stimulation on PFC activity

The present data demonstrated that dopamine’s impact on PFC activity is influenced by the behavioral state of the animal. The complex interaction between behavioral state and neuromodulation of cortex has been long recognized ^61^. Neuromodulators, including dopamine, have been shown to impact arousal, sleep wake cycle, and movement^40, 61-63^. Consistent with this literature, we observed that dopamine stimulation caused behavioral activation. But how does the modulation of behavioral state itself impact the effect of dopamine on cortical networks? We hypothesize that movement/arousal and phasic dopamine activation by either cognitive/internal or external stimuli may interact synergistically. At baseline, animal movement may be initiated by the activation of the brainstem which may then activate many neuromodulator systems, including dopamine ^61^. This may result in the activation of cortical networks via diffuse projection of these neuromodulators throughout the cortex. When dopamine neurons fire in a sustained phasic pattern, the substantial increase in movement may enhance cortical gamma and population activity. In addition, the synergistic effect of movement on dopamine activation may result in a much greater impact of phasic dopamine release on cortical activity compared to phasic dopamine activity in the absence of movement. This explains the strong enhancement of gamma and population response by sustained phasic activation. Factoring out of solely movement-related cortical activity changes can also explain the partially reduced effect of sustained phasic stimulation on cortical network response when behavioral state is controlled for.

Burst VTA dopamine activation elicited a similar increase in PFC [DA]_o_ compared to sustained phasic activation but did not generate strong behavioral activation. It also produced a minimal change in cortical network activity between baseline and stimulation periods. Even though phasic dopamine release occurs in PFC during prolonged burst dopamine firing, the lack of amplification of dopamine response by behavioral activation may prevent a substantial modulation of cortical activity. An additional possible mechanism is that dopamine receptors in PFC are differently affected by these patterns of dopamine cell activity. Postsynaptic and behavioral effects of dopamine in the PFC are primarily mediated by dopamine D1-like receptors ^20, 64, 65^. Importantly, this class of receptors rapidly desensitizes after exposure to endogenous dopamine or D1 agonists by both internalization ^66^ and dephosphorylation ^67^. A potential mechanism may be that burst activation of dopamine neurons produces a rapid bolus delivery of dopamine to D1 receptors that desensitizes these receptors more effectively than the more uniform pattern of dopamine delivery after sustained phasic activation. The combination of low levels of behavioral activation and potential D1 sensitization during fast burst dopamine firing may result in the observed weak modulation of gamma and population activity in PFC. The influence of burst mode of dopamine activation on cortical activity emerges when animals become behaviorally active. This observation can be explained by the hypothesis that the synergistic interaction between behavioral activation and dopamine response outweighs the effect of desensitization of D1 receptors. Such mechanisms would allow for a high degree of flexibility on the part of the PFC neurons to modulate their response to different pattern of dopamine neuron activation by dynamically regulating the number of dopamine receptors and by adjusting to the current behavioral state of the animal.

In conclusion, we find that dopamine modulation of PFC is governed by the pattern of dopamine neuron activation and animal’s behavioral state. Phasic activation of dopamine neurons causes a weak direct response on the discharge rate of individual neurons but causes pronounced modulation of population response in neuronal state space and of LFP gamma oscillations that are sustained for seconds to minutes after the onset of phasic dopamine activation. These results indicate that dopamine enables PFC to compute and generate spatiotemporally diverse and specialized outputs that are not a direct/linear function of the input. The multiple modes of influencing PFC computational capacities are relevant for understanding dopamine’s impact on PFC-dependent affective and cognitive functions and the deficits associated with a multitude of dopamine-related brain illnesses such as schizophrenia, addictive disorders, and ADHD.

## Methods

### Subjects

Adult male transgenic Long Evans rats expressing Cre recombinase under the control of tyrosine hydroxylase promoter (*Th::Cre*) and wild-type littermates were used. Rats were housed in a 12-hour reverse light/dark cycle with lights on at 7 pm. Experiments were approved by and conducted according to the ethical guidelines of the Institutional Animal Care and Use Committee at the University of Pittsburgh.

### Stereotaxic surgery, virus infusion, and cranial implantation

Cre-inducible recombinant adeno-associated viral (AAV) viral vector constructs containing the gene encoding ChR2 (AAV5-Ef1α-DIO-hChR2-eYFP) were obtained from the University of North Carolina Vector Core (Chapel Hill, NC, USA). Under isoflurane anesthesia, 1 µL injections were made at each of the four unilateral VTA sites (AP = −5.0 and −6.0 mm, ML = 0.6 – 0.7 mm, DV = −7.0 and −8.2 mm) at a rate of 0.1 µL□min^−1^ using a microsyringe (Hamilton Co., Reno, NV, USA) and a pump (World Precision Instruments, Sarasota, FL). Either in the same surgery or a few weeks after the viral infusion surgery, an optical fiber or an optrode was implanted in VTA unilaterally while an electrode or a microdialysis cannula was implanted in the ipsilateral mPFC. Optical fibers with metal cannulae (200 µm core diameter, 0.22 NA; Doric Lenses, Quebec, Quebec, Canada) were implanted in the VTA at the following coordinates: AP = −5.4 – (−5.8) mm, ML = 0.6 - 0.7 mm, DV = −7.0 mm. Optrodes were custom-built in house by gluing an electrode array, consisting of 8 Teflon-insulated stainless steel wires, to an optical fiber; the wires extended below the optical fiber termination by ~ 500 um. The optrodes were implanted in the VTA at the following coordinates AP= −5.4 – (−5.8) mm, ML = 0.6 - 0.7 mm, DV = −7.7 mm.

An electrode array consisting of 8 Teflon-insulated stainless-steel wires (NB Labs, Denison, TX) was implanted in the mPFC at the following coordinates: AP = 3.0 mm, ML = 0.7 mm, and DV = −3.3 mm. Microdialysis guide cannulae (CMA Microdialysis, Holliston, MA) were implanted at the following coordinates: AP = 3.0 mm, ML = 0.8 mm, and DV = −1.5 mm. All coordinates are given in mm relative to Bregma, and DV coordinates are relative to the brain surface; coordinates were adjusted for individual surgeries. Experiments were conducted at least 4 weeks from viral infusion surgeries.

### Histology and immunohistochemistry

At the end of the experiment, animals were euthanized with an intraperitoneal injection of chloral hydrate (400 mg/kg), transcardially perfused with 0.9% saline and 4% paraformaldehyde (PFA) (diluted from 32% PFA solution from Electron Microscopy Sciences, Hatfield, PA, USA) and decapitated. Brains were fixed in 4% PFA overnight at 4°C and then transferred to a cryoprotectant consisting of 20% sucrose. Brains were sectioned at 35 µm to get coronal slices of the VTA, and optical fiber/optrode placements in VTA were verified by staining slices with cresyl-violet. To examine the expression of ChR2-eYFP in dopamine neurons of VTA, VTA coronal slices were treated immunohistochemically as previously described ^25^. To verify electrode and microdialysis cannula placements, 60 µm coronal slices of the mPFC were stained with cresyl-violet and observed under a light microscope.

### Optical stimulation of VTA

For VTA stimulation, a patch cord (200-µm core diameter, 0.22 NA; Doric Lenses), attached to a 473-nm blue laser diode (OEM Laser Systems, Midvale, UT) and controlled by a Master-8 pulse generator (A.M.P.I., Jerusalem, Israel), was connected to the implanted optical fiber with a zirconia sleeve (Doric Lenses). Laser output was measured at the optical fiber tip before each experiment with a broadband power meter (Thor Labs, Newton, NJ). VTA was stimulated with a repeated phasic stimulation sequence that utilized slow burst, fast burst, and sustained phasic protocols (Figure 1C). In the slow burst protocol, VTA was stimulated with 200 ms bursts (20 Hz, 5 ms pulse width, 4 pulses) repeated every 500 ms for 10 min. In the fast burst protocol, VTA was stimulated with 200 ms bursts (100 Hz, 5 ms pulse width, 20 pulses) repeated every 500 ms for 10 min. In the sustained phasic protocol, VTA was stimulated continuously for 5 s (20 Hz, 5 ms pulse width, 100 pulses), and the phasic trains were repeated every 10 s for 20 min. The sustained phasic stimulation was delivered for a longer period compared to the burst protocols to acquire a sufficient number of transient trials (each 10-s period was considered a trial, ~120 total trials in 20 min). On the other hand, fast and slow burst stimulation protocols allowed for a greater number of transient trials (~1200 in 10 min). Light output was measured to be 5 – 15 mW in *Th::Cre* and 9 – 15 mW in wild-type electrophysiology sessions. In microdialysis experiments, light output was measured to be ~10 mW in all *Th::Cre* and wild-type sessions.

### Microdialysis Procedures

Rats (*Th::Cre*: n = 4 and wild-type: n = 3) were quickly and lightly anesthetized with isoflurane for insertion of the microdialysis probe (membrane length = 3 mm, CMA Microdialysis) into the guide cannulae without tearing the probe membrane. Probes were perfused with Ringer's solution (in mM: 37.0 NaCl, 0.7 KCl, 0.25MgCl_2_, and 3.0 CaCl_2_) at a flow rate of 2.0 μL/min during sample collection. Dialysate samples were collected every 20 min and immediately injected into an HPLC system for electrochemical detection of dopamine as described before ^68^. After establishing at least three stable baseline samples, VTA was stimulated with either the sustained phasic or the fast burst protocol for 20 min. Animals were allowed to recover from isoflurane inhalation for at least 90 min from probe insertion before collection of the first baseline sample used in data analysis. Each animal was run over multiple sessions (with at least 24 hours between sessions), but only one session per protocol per animal was used for data analysis. Other sessions were removed because of leaking microdialysis probes and low sample volume, inability to separate dopamine peaks in the chromatogram, or patch cord breakage.

### Electrophysiology Procedures

For awake recordings, the implanted electrodes were connected to a unity-gain junction field effect transistor headstage and lightweight cabling, which passed through a commutator and enabled rats to move freely (Plexon, Dallas, TX). Signals were digitized at 40 kHz sampling rate, and then band pass filtered at 0.5 – 200 Hz (or 125 Hz in some cases) for LFP channels, at 300 Hz – 8kHz for mPFC spike channels and at 150 Hz – 8kHz for VTA spike channels. LFPs were amplified at 500X gain and downsampled to 1 kHz via an OmniPlex acquisition system (Plexon). The signal was referenced against a ground that was placed in the skull above the cerebellum.

To acquire neural spikes, continuous spike channel signals were thresholded using a threshold that exceeded baseline noise by at least 3 SD. Voltage waveforms that crossed the threshold were isolated into clusters using Plexon offline sorter (Plexon); clusters that were well isolated from other unit clusters and from noise were designated as single units (SUA) and unit clusters that did not separate well from each other were classified as multi-units (MUA).

### General recording session procedures

#### mPFC recording

Recordings were conducted in freely moving rats (*Th::Cre*: n = 17 and wild-type: n = 4). Rats were habituated to the recording chamber (12”×10“×12“) for 1-2 days prior to the first recording session. Each recording session consisted of 30 minutes of baseline recording, after which light pulses were delivered to the VTA using one of the repeated phasic stimulation protocols described above. Recording continued for 60 minutes after stimulation onset. Only one stimulation protocol was delivered per session, and the order of protocols across sessions was counterbalanced across animals (minimum period of 24 hours between sessions). Most animals were presented with a particular stimulation protocol only once. A few animals received multiple sessions of the same stimulation protocol because some sessions had to be removed due to suspected fiber breakage, instability of the laser power, and/or other experimental issues.

#### VTA optrode recording

A subset of *Th::Cre* rats (n = 4) used in the awake mPFC recordings were also implanted with optrodes in VTA to examine responses of VTA units to laser stimulation. Recordings in the VTA were conducted either in the same session as the mPFC recordings or in a separate session. In a few sessions for which mPFC recordings were not used for analysis, laser output was adjusted on-line (between 1-18 mW); power was increased to elicit activation in VTA units if no response was detected or reduced if multi-unit response or large field response to stimulation prevented isolation of single units.

#### Movement tracking

Movement activity was tracked in most of the freely moving recording sessions via an infrared activity monitor (Coulbourn, Holliston, MA).

### Data Analysis

Data analysis was conducted with custom scripts in Matlab (Mathworks) and SPSS statistical software (IBM). Hyunh-Feldt correction was applied to repeated measures ANOVAs to correct for any sphericity violation. Multiple comparison correction was applied for more than 3 post-hoc comparisons. Parametric or non-parametric statistical tests were used depending on the data.

#### Microdialysis data

Dopamine concentration (fmol/μL) for each sample was expressed as a percentage of average baseline dopamine concentration (baseline constituted three samples immediately before stimulation). For statistical analysis, % values were first log transformed to reduce the large variability in dopamine concentration increases upon stimulation. Repeated measures ANOVA was conducted on log transformed values, and one sample tests against baseline average were performed for post-hoc analysis; one tailed t-tests were used to test the hypothesis that stimulation elicited an increase in dopamine levels. A two way repeated measures ANOVA was also run to compare mPFC extracellular dopamine responses to VTA stimulation between *Th::Cre* and wild-type rats and between stimulation protocols in *Th::Cre* rats.

#### VTA optrode data

To assess the responses of VTA single units to optogenetic VTA stimulation, perievent histograms and rasters were examined. For each unit, the probability of spiking at different latencies from the time of a single blue laser pulse delivery was also calculated.

#### mPFC unit classification

Single units recorded in mPFC were classified as regular-spiking (RS) or fast-spiking (FS) based on their firing rate at baseline (before stimulation) and spike waveform characteristics ^69^. As FS units fire at higher frequencies and exhibit narrow spike waveform widths, units were identified as FS if firing rate >=10 Hz, valley width at half height < = 0.3 ms, and peak to valley width < = 0.3 ms. Based on this criterion, 7 units were classified as FS and the rest as RS units.

#### Movement analysis

Based on digital movement scores and manual recordings of animal behavior, 3 min epochs in baseline, stimulation, and post-stimulation periods during which animals were continuously moving at comparable levels were manually identified. For *Th::Cre* rats, two epochs (stim 1 and stim 2) were extracted during the stimulation period. As wild-type rats were less mobile, only one movement epoch was extracted during the stimulation period; to keep the number of epochs for statistical comparison similar to *Th::Cre* rats, two movement epochs were isolated in the post-stimulation period. If at least two 3-min movement epochs in *Th::Cre* sessions and one 3-min epoch in wild-type sessions couldn’t be identified during the stimulation period, those sessions were removed from analyses that involved movement epochs only. As this method resulted in the exclusion of many wild-type sessions, data from all three stimulation protocols were combined for wild-type rats (n = 6 sessions from two rats) to increase the sample size. All extracted post-stimulation epochs occurred at > 15 min from offset of stimulation sequence.

#### Transient unit activity around each phasic stimulation train

***Individual unit response**.* Transient responses of prefrontal units to phasic VTA stimulation were examined by binning spike counts (bin size = 1s for sustained phasic protocol and bin size = 0.05 s for fast and slow burst protocols) and calculating firing rates. Independent t-tests were conducted between each bin’s firing rate against mean baseline firing rate, and units were deemed to be significantly modulated if at least one post-stimulation bin was significantly different from baseline at p < = 0.001. P < = 0.001 criterion was more conservative than Bonferroni correction, but this criterion prevented false positives based on examination of rasters and histograms. Color maps were used to visualize responses of all units; values in the color map indicated changes in firing rate from baseline represented by t-values.

***Population response.*** To examine changes in population activity patterns, a distance-based similarity measured was used. For this measure, each population state at a particular time was comprised of spike counts of N-units in the N-dimensional space, forming a spike count vector. We then calculated Euclidean distance between population states during stimulation and baseline to assess whether the population states diverged over time from baseline as a result of phasic VTA dopamine activity.

Units with firing rates < 0.1 Hz were removed from this analysis. To get pseudo-baseline trials, the baseline period was binned into consecutive trials of the same duration as stimulation trials (10 s for sustained phasic stimulation and 0.5 s for fast and slow burst stimulation). Within each trial, spike counts were calculated in bins (bin size = 2 s for sustained phasic stimulation and 0.1 s for fast and slow burst stimulation). Spike counts for each bin were pooled across all recorded single and multi-units (not just simultaneously recorded) to get pseudo-population spike count vectors. Then, Euclidean distances were calculated between spike count vectors from pseudo-baseline trials and stimulation trials in each bin. One tailed two sample t-tests (Bonferroni corrected) between distances in the first bin and all other bins were conducted to assess if the population activity diverged over time because of VTA stimulation (which would correspond to increased distances, thus right t-tailed t-tests were used).

#### Sustained unit activity during repeated phasic VTA stimulation

***Individual unit response.*** Unit spike counts were binned into 120 s to get firing rates. Firing rates were Z-score normalized against baseline, and units were determined to be significantly activated or inhibited if firing rates post-stimulation were respectively greater or less than 99% confidence interval (corresponding to Z-scores > or < 1.96) of baseline in 3 consecutive bins ^70^. Z-values rather than t-values were used to get confidence intervals and assess significance as there were no trials in this analysis, and each unit’s activity was assessed over the course of a session. Thus, firing rates across baseline bins were used to calculate baseline standard deviation for obtaining Z-values; t-values would, in this case, be highly influenced by the number of baseline bins. Proportions of units activated/inhibited by stimulation were calculated, and a logistic regression model with group and stimulation protocol as predictors (interaction term excluded) was fit to determine difference in the proportion of unit modulation between *Th::Cre* and wild-type rats and across stimulation protocols.

***Population response.*** To assess sustained population responses to prolonged VTA stimulation, Euclidean distances of spike count vectors were calculated between baseline and stimulation. Spike counts were pooled across all recorded single and multi-units to get pseudo population response vectors. Units with firing rates < 0.1 Hz were removed. The entire session was divided into epochs of 5 min duration, resulting in 3 baseline epochs (−15 to −10 min, −10 to −5 min, and −5 to 0 min). Each epoch was further binned into 10 s consecutive trials (total 30 trials). All pairwise distances of spike count vectors were calculated between each trial of an epoch and all trials of baseline epoch 1 and then averaged to yield one distance value per trial. To determine whether the population response significantly diverged from baseline after stimulation, distances for each epoch were compared against distances for baseline epoch 2 using right-tailed two sample Bonferroni corrected t-tests.

When population activity was analyzed in movement epochs only, each epoch was divided into 10 s consecutive trials (total: 18 trials). Spike count vector distances between each trial of an epoch (stimulation or post-stimulation) and all trials of the baseline epoch were averaged to yield a mean pairwise distance value per trial. To assess the divergence of population activity upon stimulation, these mean pairwise distance values were compared between stimulation and post-stimulation epochs using one-tailed Bonferroni-corrected two sample t-tests. For visualization of high dimensional population activity during different epochs, spike count vectors were analyzed with PCA, and scores were projected onto any two of the first three dimensions of the PCA space.

#### Transient LFP response to each phasic stimulation train

Transient LFP data were analyzed using the continuous wavelet transformation (CWT, with complex Morlet wavelets) via NDTools toolbox (http://spot.colorado.edu/~gilley/index.html). A CWT rather than fast-fourier transform (FFT) was used for transient analysis of LFP power because the former method is especially suited to detect transient changes in LFPs, and FFT frequency resolution depends on the bin size (which would be around 0.1 s for fast and slow burst stimulation trials; this bin size would highly reduce the frequency resolution and wouldn’t permit examination of power in low frequencies). LFPs were detrended, artifacts were removed, and CWT was applied to LFP from each stimulation trial (that was 10 s in duration for sustained phasic stimulation and 0.5 s for fast and slow burst stimulation). LFP power at each frequency and time point was averaged across trials and Z-score normalized to the total power within each frequency. Frequencies were binned into several bands: delta (1 - 4 Hz), low theta (4.5 – 8 Hz), high theta (8 - 13 Hz), beta (14 – 30 Hz), low gamma (30 – 55 Hz), and high gamma (55 – 100 Hz). For statistical analysis, normalized LFP power within each frequency band was divided into 1 s bins (for sustained phasic stimulation, trial window = [-2 s 8 s]) or 0.05 s bins (for fast and slow burst stimulation, trial window = [-0.1 s 0.4 s]), and repeated measures ANOVAs followed by Bonferroni corrected post-hoc paired t-tests against baseline (bin 1) were used to assess changes in LFP power within each frequency band over time. To obtain scatter plots of baseline power versus stimulation power, baseline power was determined by averaging across two baseline bins and the last post-stimulation bin while stimulation power was calculated as an average of the first three bins from stimulation onset.

#### Sustained LFP response to repeated phasic VTA stimulation

Even though LFPs were recorded for 30 min of baseline and 60 min post stimulation onset, we observed that the early baseline period prior to −10 min (from stimulation onset) in all three paradigms was characterized by long periods of enhanced lower frequency activity and reduced high frequency oscillations, which is indicative of a drowsy or sleepy rat ^32^. Similarly, the post-stimulation period beyond 20 min from stimulation onset showed an increase in the power of lower frequencies and decrease in higher frequencies (Figure 9). Based on these LFP observations and manual observations of rat behavior, the whole session LFP statistical analyses depicted in Figure 9 were limited to a baseline period of 10 min immediately prior to stimulation onset and a post-stimulation period that lasted from stimulation onset to 5 min from stimulation offset (−10 min to + 25 min for sustained phasic stimulation sessions and −10 min to +15 min for fast and slow burst stimulation sessions).

LFP was detrended, artifacts were removed, and data were analyzed using the Chronux toolbox (http://www.chronux.org). LFP spectral power was calculated using a multi-taper FFT with a sliding time window of 4 s in 2 s steps. A standard multi-taper approach was used that applied the 15 leading tapers to each window (time bandwidth product = 8). Each session’s spectrogram was Z-score normalized to baseline, and Z-scored LFP power for each frequency band was divided into 5 min bins. A repeated measures ANOVA (followed by post-hoc Bonferroni corrected paired t-tests against baseline) was conducted to detect any difference in spectral power across time.

For the analysis of LFPs in moving epochs, a multi-taper FFT was conducted with the same parameters as in the whole session analysis. Each epoch’s spectrogram was Z-score normalized against the baseline and post-stimulation epochs (post-stimulation 2 for wild-type). Z-scored LFP power values across all time points within an epoch were averaged for each frequency band, and the effect of stimulation was assessed by a repeated measures ANOVA followed by paired t-tests.

#### Assessment of phase locking of spikes to LFPs

To correct for phase lags in lower frequencies, LFPs were first aligned using FPAlign (Plexon). Signals were then band-pass filtered using eegfilt.m (using the least squares finite infinite response filter) function from the EEGLab toolbox (https://sccn.ucsd.edu/eeglab/index.php). Band-pass filtered LFPs were Hilbert transformed to extract instantaneous phases, and a circular distribution of phases for each unit was obtained by extracting phases corresponding to each spike. For the entire session analysis, LFPs and unit spike counts were divided into 5 min bins. Single units and multi-units were pooled, and only units that emitted at least 100 spikes within each bin were included. The circular statistic toolbox ^71^ was used to perform Rayleigh’s Z-test for circular uniformity and determine if units were significantly phase locked to specific bands within each time bin. Proportions of significantly phase locked units were compared across time bins using Fisher’s exact tests. To determine the strength of phase-locking to each frequency band, phase locking values (PLVs) were computed for all units. PLVs correspond to the mean resultant length (MRL) of phase angles. In short, MRL is calculated as the modulus of the sum of unit vectors representing instantaneous LFP phases corresponding to each spike divided by the number of spikes ^72^. To assess a change in the strength of phase-locking over time, PLVs for all units (not just significantly phase locked units) were compared using Friedman’s ANOVA followed by Bonferroni-corrected signed rank tests.

#### Assessment of phase amplitude coupling in LFPs

LFPs were band pass filtered using the EEG toolbox and Hilbert transformed within each band to extract instantaneous phases and amplitudes. Phase amplitude coupling between the phase of low frequencies (delta, low theta, and high theta) and the amplitude of higher frequencies (low gamma and high gamma) was assessed by calculating modulation indices (MI). MI was calculated using the get_mi.m function in the cross frequency toolbox (available from http://accl.psy.vanderbilt.edu/resources/analysis-tools/cross-frequency-interactions/) that is based on Tort et al. (2010)’s method ^73^. MI depends on the Kullback-Leibler distance between an empirical distribution containing amplitudes of one signal (high frequency in this study) over phase bins of another signal (low frequency in this study) and a uniform distribution.

MI values for each frequency pair were calculated in 5 min bins and normalized for each animal by dividing by the average MI value in the session. Differences in MI values over time were assessed via a Friedman’s ANOVA followed by Bonferroni corrected signed rank tests. To rule out spurious effects, surrogate sets were built by randomly partitioning LFP time series into two slices and rearranging the order of the slices; 100 surrogate sets were built and the MIs for all surrogate time series were averaged to get a surrogate MI value per animal. In the movement epoch analysis, MI values were normalized by the average MI value across epochs and were assessed for effects of stimulation using a Friedman’s ANOVA followed by signed rank tests.

## Acknowledgements

This work was supported by the National Institute of Mental Health grant MH48404 and MNTP fellowship at the University of Pittsburgh. Authors are grateful for helpful discussions with Dr. Marlene Cohen.

